# Structural and biochemical basis for cannabinoid cyclase activity in marine bacterial flavoenzymes

**DOI:** 10.1101/2025.10.07.680811

**Authors:** Anna C. Love, Harshverdhan Sirohi, Felix M. Hubert, Ying-Chieh Kao, Daniel E. Quinnell, Ryan Gappy, Marissa Sheehy, Jonathan Hsu, Adrian Lee, Liana Zangwill, Bruce A. Palfey, Geoffrey Chang, Bradley S. Moore

**Affiliations:** Center for Marine Biotechnology and Biomedicine, Scripps Institution of Oceanography, University of California, San Diego, La Jolla, CA 92093-0202, United States of America; Skaggs School of Pharmacy and Pharmaceutical Sciences, 9500 Gilman Drive 0754, University of California, San Diego, La Jolla, CA 92093, United States of America; Department of Pharmacology, School of Medicine, 9500 Gilman Drive 0754, University of California, San Diego, La Jolla, CA 92093, United States of America; Department of Biological Chemistry, University of Michigan, 1150 West Medical Center Drive, Ann Arbor, MI, 48109-0606, Michigan, USA

## Abstract

The marine bacterial flavoenzymes Clz9 and Tcz9 can process cannabigerolic acid (CBGA) to the minor cannabinoid, cannabichromenic acid (CBCA), however, the mechanistic details of this extrinsic transformation are still obscure. Here, we report a thorough analysis of CBCA-formation by Clz9 and Tcz9 through high-resolution crystallographic characterization, biochemical analysis, and spectroscopic interrogation. Our work reveals that Clz9 and Tcz9 use different biochemical mechanisms from *Cannabis* cyclases and each other in their production of CBCA. Collection of a high-resolution substrate-bound structure, the first for any cannabinoid cyclase, provides key insights into how active site architecture affects substrate binding and stereoselectivity. Engineering approaches improve the stereoselectivity of CBCA formation by Clz9 and Tcz9, providing access to (*R*) and (*S*)-CBCA. Collectively, our work advances understanding of enzymatic cannabinoid formation and cements Clz9 and Tcz9 as two unique members of the BBE-like enzyme family with encouraging potential for biocatalytic cannabinoid production applications.

## Introduction

The berberine bridge-like (BBE-like) enzymes comprise a subfamily of vanillyl-alcohol oxidase (VAO) / *p*-cresol methylhydroxylase (PCMH) flavin-dependent enzymes that perform diverse oxidative reactions.^1^ BBE-like enzymes are characterized by a unique domain (pfam: 08031) involving a Y/FxN motif near the C-terminus that is believed to shape and structure the oxygen and flavin binding pockets.^2^ Most members of the BBE-like enzyme family also have bicovalent linkages to their flavin cofactor through conserved histidine and cysteine residues, which raises the redox potential of this family by >300 mV (**Figure 1A**).^3^ Thus, amongst the VAO/PCMH family, the BBE-like enzymes are distinct in their involvement in secondary metabolite biosynthesis and ability to catalyze several different kinds of common oxidation chemistries, including carbon-nitrogen bond formation, selective alcohol oxidation, monoterpene alkaloid oxidation, and *ortho*-quinone methide generation (**Figure 1B**).^4^ Despite these features, the BBE-like enzyme family remains exceptionally understudied, with only 0.2% of all pfam entries accompanied by structural data.^5^ Of these, even fewer have been thoroughly biochemically characterized.

**Figure 1.**
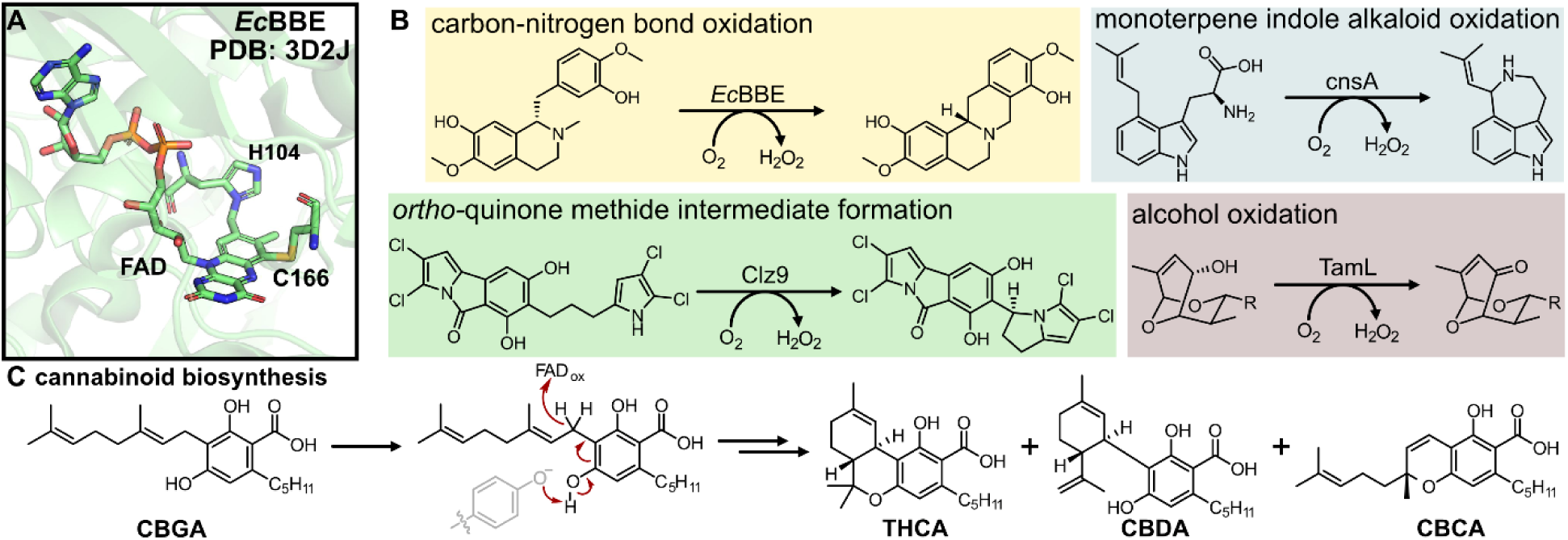
Structure and biochemical diversity of the berberine-bridge like enzyme family. (A) Bicovalent attachment of the flavin co-factor in the titular member of the BBE-like enzyme family. (B) Four general types of chemical reactivity found in the BBE-like enzyme family. (C) Mechanism of cannabinoid formation through BBE-like enzymes in *Cannabis*.

Several well-known BBE-like enzyme family members for which some structural data exists are the cannabinoid cyclization enzymes from *Cannabis sativa*, THCA synthase, CBDA synthase, and CBCA synthase.^6, 7^ To date, one crystal structure for THCA synthase has been deposited into the PDB (PDB: 3VTE),^8^ but structural data for both CBDA synthase and CBCA synthase are still lacking. It is thought that these oxidative cyclization enzymes generate *ortho*-quinone methide intermediates *en route* to product formation *via* deprotonation from conserved tyrosine residues (**Figure 1C**).^8–11^ This deprotonation initiates hydride transfer from the benzylic methylene of the geranyl chain to the N5 position of the flavin cofactor. After hydride transfer, the three enzymes enforce remarkable selectivity in the cyclization chemistry to generate three structurally distinct molecules, THCA, CBDA, and CBCA. However, the enzymes from the plant are recalcitrant to expression in bacterial hosts, likely from the signaling peptide localizing them to the glandular trichomes in native producers,^12, 13^ thus limiting their application for biotechnological production of cannabinoids to eukaryotic hosts.^14, 15^

Recently, two bacterial BBE-like enzymes that express well and with high catalytic activity in bacteria, Clz9^16^ and Tcz9^17^, were found capable of the same electrocyclization chemistry as CBCA synthase.^18^ Clz9 and Tcz9 can generate CBC, CBCA and analogs from resorcinol precursors with greater substrate flexibility than observed from the plant enzymes. While the cyclization enzymes from *Cannabis sativa* require a carboxylic acid for activity (e.g. CBGA),^8^ Clz9 and Tcz9 were capable of processing non-acid substrates, like CBG, directly into their more brain-penetrant products (e.g. CBC), making them a desirable route to directly access the more bioactive, non-acid products. Notably, this cannabinoid cyclization activity was relatively unique amongst a panel of bacterial BBE-like enzymes tested based on sequence similarity,^18^ underscoring the need to better understand the mechanism of this transformation.

Here, we report high-resolution structures of Clz9 and Tcz9 together with mechanistic biochemical analyses to evaluate the basis for their unique CBCA-production abilities. We established that Clz9 and Tcz9 are members of the BBE-like enzyme and used a structure of active Tcz9 enzyme with CBGA substrate bound to begin dissecting the mechanism for CBCA formation, finding that they operate through different pathways based on the structure and function data. These pathways were interrogated spectroscopically, which show major differences in rate and ability to oxidize substrate analogs. Finally, we used insights from both the structures and mechanistic analyses to guide rational improvement of the stereochemical outcome for CBCA-formation through both substrate and enzyme level engineering. Collectively, our study reveals new structural and chemical insights into two unique bacterial BBE-like enzymes, an understudied family of flavoproteins with promising potential for biocatalytic application.

## Results

### Clz9 and Tcz9 are members of the BBE-like enzyme family

To establish Clz9 and Tcz9 as members of the BBE-like enzyme family, we determined the crystal structures of the enzymes with flavin cofactor bound at 1.95 and 1.56 Å, respectively, *via* molecular replacement (**Figure 2**, **Table S1**). Structural analysis revealed that both enzymes contain the VAO-fold,^19^ bicovalent attachment of the flavin cofactor,^3, 20, 21^ and the Y/FxN motif at the C-terminus.^22^ The VAO-fold is characterized by seven antiparallel β-sheet fold associated with flavin-binding, and several alpha helical loops (**Figures 2A, S1**).^19^ The namesake of the BBE-like enzyme family, responsible for constructing the C–C berberine bridge bond to generate (*S*)-scoulerine from (*S*)-reticuline in the California poppy plant, has both an 8α-*N*^1^-histidyl–FAD linkage through His104, and a 6-*S*-cysteinyl–FAD linkage through Cys166.^23^ While some members of the BBE-like enzyme family have lost one (e.g. EncM,^24^ MaDA^25^) or both^26^ of the covalent bonds, Clz9 and Tcz9 maintain both attachments *via* His99/Cys157 or His67/Cys125, respectively (**Figures 2B-D**). In the apo models, both Tcz9 and Clz9 show a high degree of similarity and superimpose with overall RMS of (0.681 Å).

**Figure 2.**
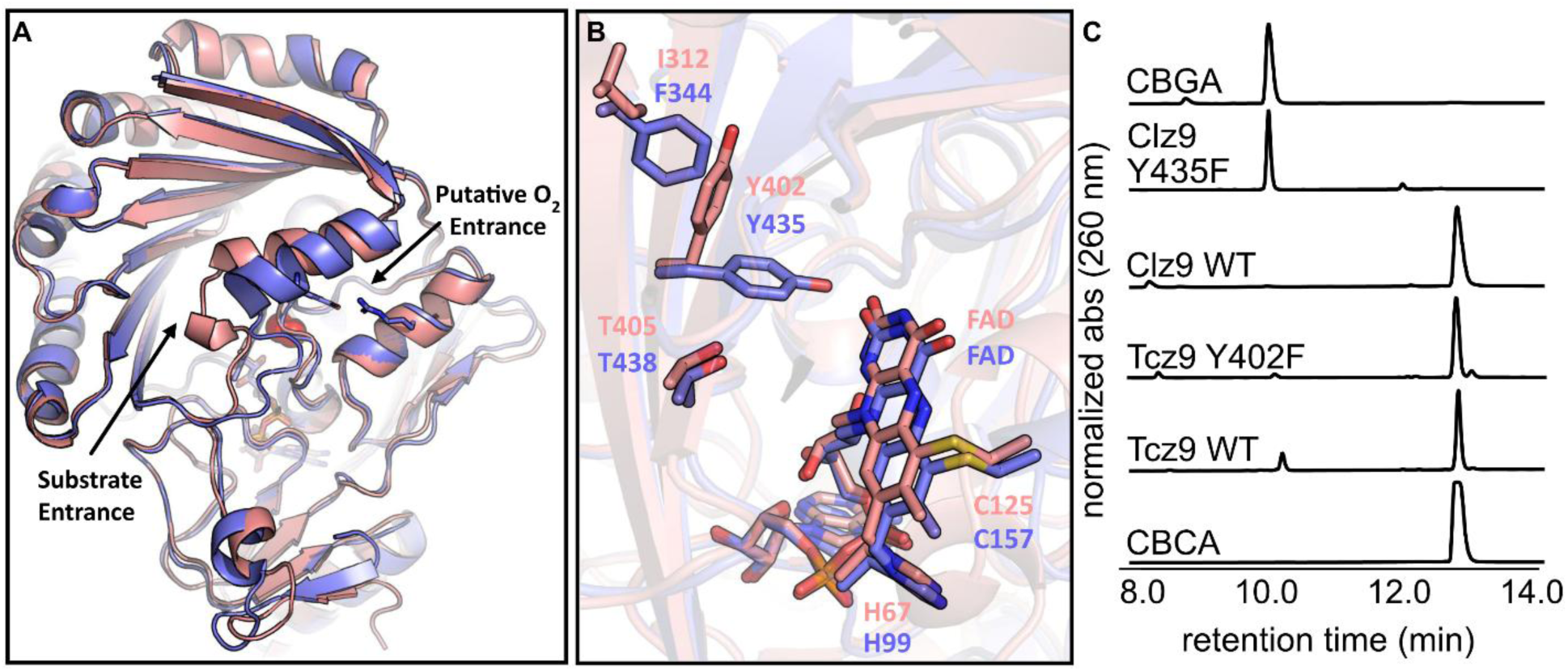
Structural characterization of Clz9 and Tcz9. (A) Superposition of Tcz9 (pink) and Clz9 (purple) structures. The entrance/exit for substrate and the position of the putative “oxyanion hole” are pointed out with arrows. A dioxygen model (red sphere) near the N5 of the FAD is also shown. (B) Close-up of the Y/FxN motif of Tcz9 (pink) and Clz9 (purple), showing the relative positional differences in the placement of Tcz9-402 and Clz9-435 causes by Tcz9-I312 and Clz9-F344. Also shown are the covalent linkages due to conserved histidine (Tcz9-H67, Clz9-H99) and cysteine (Tcz9-C125 and Clz9-C157). (C) Biochemical validation of the differing tyrosine rotamers in Clz9 (e.g. Clz9-Y435F) and Tcz9 (e.g. Tcz9-Y402F).

The largest difference between the Clz9 and Tcz9 structures is in a helix-loop region (Tcz9 residues 284-302; Clz9 residues 316-334) that likely plays a role in the entrance of substrate and the egress of product from the active site (**Figures 2A, S2**). We speculate that this helix-loop structure could also play a role in substrate specificity and dioxygen entry. In Tcz9, this loop includes a stretch of four consecutive alanine repeats (QGEAAAAEQVR) while in Clz9, the corresponding region is different (QGEAMKATSVR). These sequence differences manifest in a significant structural difference in the loop positioning between the two enzymes, with an RMSD of 4.22 Å between Tcz9/Clz9 residues 295-305/327-337 respectively (**Figures 2A and S2**). In Clz9, E324 makes a notable salt bridge with R143 (**Figure S2**) that obstructs an opening to a tunnel leading to the active site near FAD. This salt bridge is absent in the Tcz9 structure (**Figure S2**). Other BBE-like enzymes, including the plant THCA synthase, have even bulkier loops that block this entrance (**Figure S3**).^8, 22, 23^ Interestingly, the corresponding residues in Tcz9 (Tcz9-E292 and Tcz9-N111) on this loop do not form a salt bridge and the tunnel is wide open (**Figures 2A, S2**). This opening leads to a small cavity reminiscent of the ‘oxyanion hole’ observed in certain hydrolases,^22^ and is the likely site to orient molecular oxygen in BBE-like oxidases for regeneration of oxidized flavin (**Figure S4**).

Previous comparison between two BBE-like flavoenzymes, an oxidase that reduces molecular oxygen rapidly from the plant *E. californica* (PDB 3D2H) and another slower dehydrogenase called ‘Phl p 4.0202’ (PDB 4PVE), resulted in identification of “gate-keeper” residue position which interacts with molecular oxygen.^22^ A mutation from isoleucine to valine in Phl p4 causes a 60,000x increase in dioxygen reactivity presumably by abating a steric clash with molecular oxygen in the wild type enzyme. An inverse exchange converting this position from valine to isoleucine in BBE-enzyme from *E. californica* decreased molecular oxygen reactivity by 500x through a steric blockage. In both Clz9 and Tcz9, the ‘gate-keeper’ position is in fact, an isoleucine residue (Clz9-I160, Tcz9-I128). However, it is positioned farther from the oxyanion hole (**Figure S4**) due to a small displacement of a short loop (Tcz9 residues 122-129; Clz9 residues 154-161), thus allowing molecular oxygen to better occupy the site.

Clz9 and Tcz9 also have the characteristic Y/FxN motif. When the Tyr is present in this motif, it is typically believed to create an H-bonding network involving a carbonyl of the isoalloxazine ring in flavin.^22^ However, the structure of Tcz9 demonstrates a departure from conventional BBE-like enzyme biochemistry as the specific rotamer of Tyr present in this motif is rotated 90° away from FAD (**Figure 2B, S2**, and **S5**). In Clz9, the presence of phenylalanine at position 344 likely forces the tyrosine toward FAD, while in THCA synthase a tyrosine (Y481) performs a similar function. In Tcz9, this position is occupied by an isoleucine (I312), and the reduced steric bulk allows space for the alternative tyrosine rotamer to form. In other BBE-like enzymes, like the glucooligosaccharide oxidase from *Sarocladium strictum* (PDB: 1ZR6),^27^ a similarly small, nonpolar residue is present at this same position (A323), but the tyrosine in the Y/FxN motif is pointing toward the FAD like in Clz9 and THCA synthase. Thus, having enough space is not the sole reason the alternative tyrosine rotamer may form. It is likely the H-bonding interaction between Y402 and nearby M344 is the driving force for formation of this rotamer or by steric interactions with surrounding ordered waters. To the best of our knowledge, this is the first time this rotamer of the tyrosine residue in the Y/FxN motif has been observed in BBE-like enzymes.

To further establish that the tyrosine in Tcz9 is not involved in flavin stabilization/orientation, point mutants from tyrosine to phenylalanine were generated from both Clz9 and Tcz9. As expected, the Clz9 Y435F mutant lost activity with CBGA as a substrate, while Tcz9 Y402F retained full conversion of CBGA to CBCA (**Figure 2C**). Collectively, these data establish that Clz9 and Tcz9 are members of the BBE-like enzyme family with some unique features that expands our understanding of potential BBE-like enzyme structures.

### Clz9 and Tcz9 have different mechanisms for cannabinoid cyclization

The ability for Clz9 and Tcz9 to catalyze cannabinoid cyclization chemistry is relatively unique amongst tested bacterial BBE-like enzyme family members.^18^ To better understand how Clz9 and Tcz9 carry out this chemistry, an X-ray crystallographic structure of Tcz9 with the substrate CBGA bound was generated to compare to the apo structures (**Figure 3 and S6**). Excitingly, when wild type Tcz9 crystals were soaked with CBGA prior to X-ray diffraction analysis, the substrate was “caught in the act” despite using catalytically active enzyme. We believe that this Tcz9-CBGA complexed structure represents a locked state in the context of the crystal. We found that there were two conformers of CBGA present based on the electron density map, which were present in roughly equal quantities (**Figure 3, S6**), displacing ordered water in the apo structure of Tcz9 (**Figure S6**). The substrate-bound structure was examined for polar contacts between active site residues and substrate (**Figure 3**). While the formal mechanism for cannabinoid cyclization in plant enzymes has not yet been established, the proposed mechanism involves deprotonation by a tyrosine residue conserved amongst all three cyclases from *Cannabis sativa* (**Figure S7A**). Recently, another mechanism was proposed that differed slightly in the order of operations, but still relied on initial deprotonation with the same tyrosine residue (**Figure S7B**).^28^ This tyrosine residue, however, is not conserved in Clz9 and Tcz9, and is instead replaced with threonine (Clz9-T438 and Tcz9-T405). When mutated to a valine, both Clz9 and Tcz9 retained full conversion of CBGA to CBCA, indicating that a different deprotonating residue was responsible (**Figure S7C**).

**Figure 3.**
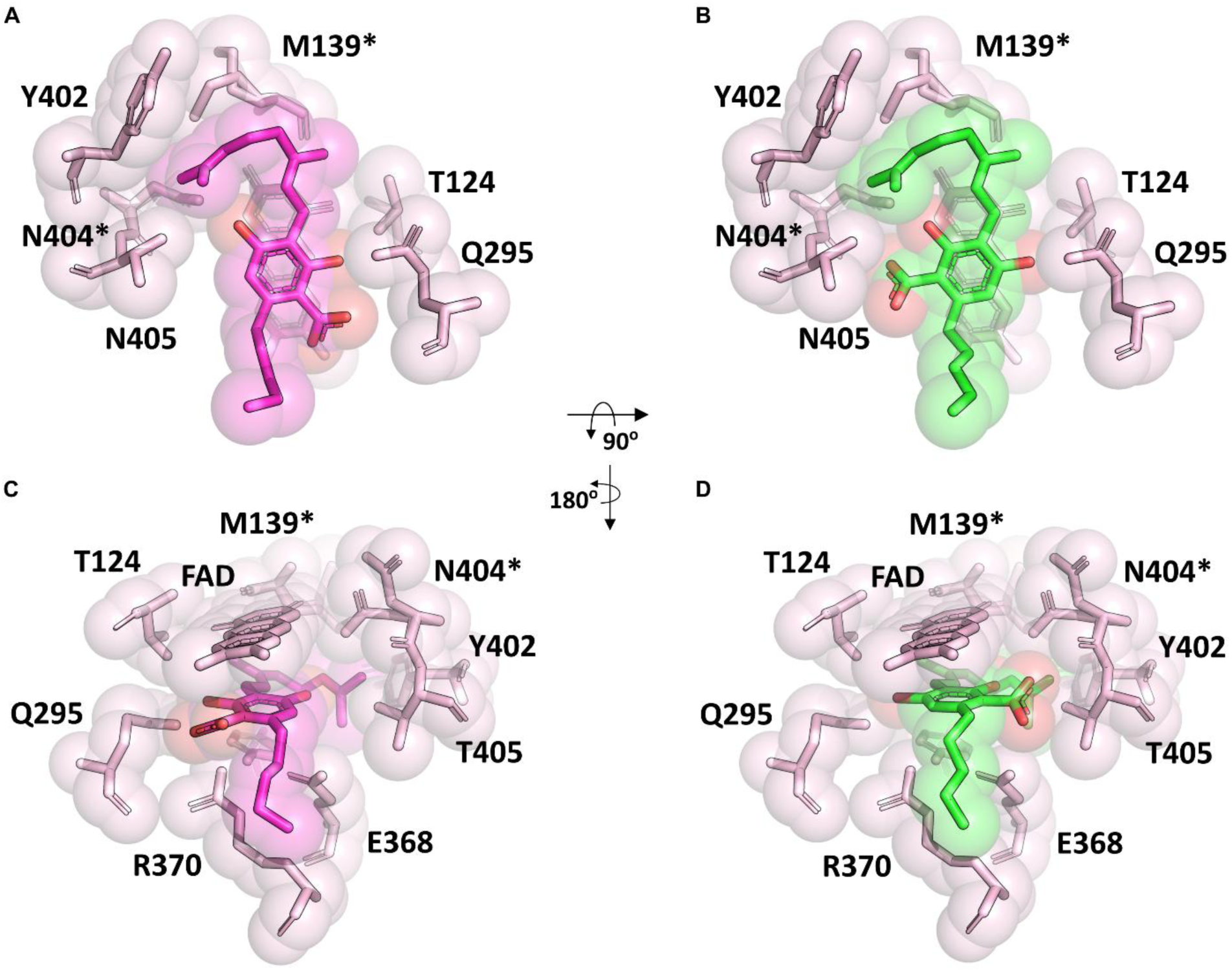
Tcz9 in complex with CBGA. Views of the Tcz9 enzyme active site with CBGA bound looking (A-B) perpendicular to and (C-D) in the plane of flavin. There are two conformations of CBGA (colored purple and green) in this crystal form due to the pseudo symmetry of the CBGA molecule with the differences in the carboxylic acid position. Each alternative conformation of CBGA captures, colored purple (A, C) and green (B, D) are shown separately. Tcz9 residues surrounding the CBGA are indicated. Residues denoted with an asterisk have alternative conformations in the electron density map. The approximate rotational transformations relating the two views are indicated.

Also conserved amongst the three plant enzymes is another tyrosine residue that is proposed to form a H-bonding interaction with the non-reactive phenol in the resorcinol substrate.^8^ This tyrosine residue is conserved in both Clz9 (Y374) and Tcz9 (Y342), however, in the substrate bound structure of Tcz9-CBGA, Y342 does not form a hydrogen bond with CBGA and, in this trapped state, is also not in close enough proximity for deprotonation (**Figure 4C**). Thus, we hypothesized this tyrosine residue was likely not involved in the deprotonation of CBGA. In Clz9, Y374 is pointed on a slightly different trajectory than in Tcz9, hinting that perhaps it plays a different role in Clz9. Thus, tyrosine to phenylalanine mutants were generated for both enzymes and mutants incubated with CBGA. As expected, the Tcz9-Y342F mutant retained full conversion to CBCA, confirming that it is not involved in the mechanism of CBCA formation in Tcz9 (**Figure 4B**). However, the Clz9-Y374F mutant had a >95% reduction in activity compared to wild type enzyme (**Figure 4B**). This divergence in behavior suggests that Clz9 and Tcz9 use different mechanisms to perform cannabinoid cyclization chemistry, with Clz9 potentially using Y374 to “activate” the substrate *via* phenol deprotonation, initiating hydride transfer.

**Figure 4.**
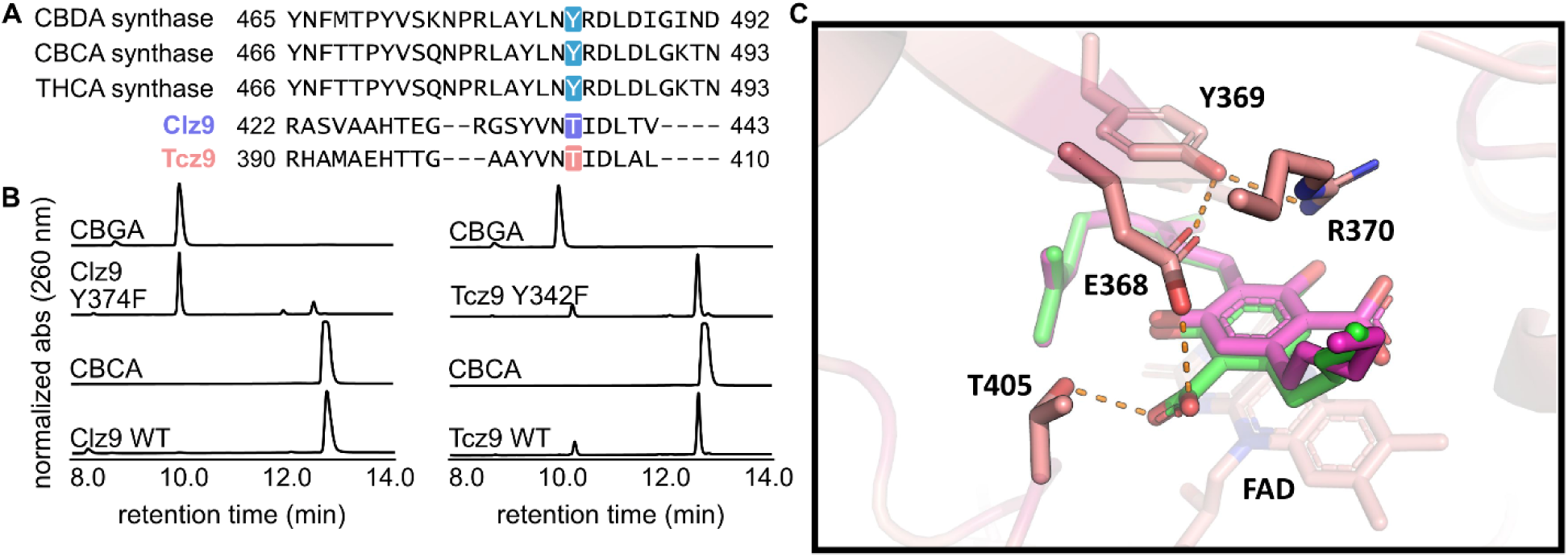
Clz9 and Tcz9 follow different mechanisms for CBCA formation. (A) MAFFT sequence alignment showing how the catalytic tyrosine residue in the plant cannabinoid synthases is not conserved in Clz9 and Tcz9. (B) Mutagenesis revealed that Clz9-Y374 was responsible for substrate activation *via* deprotonation in Clz9, but the homologous position in Tcz9 (e.g. Tcz9-Y342) did not carry out the same function. (C) CBGA-bound Tcz9 structure with hydrogen binding distances (<3.3 Å distance cutoff) are indicated as dashed lines.

### Tcz9 is able to directly perform hydride abstraction

Given the lack of polar residues in Tcz9 in the vicinity and H-bonding distance of the substrate, it was clear that a unique mechanism governed cannabinoid cyclization in Tcz9. There are relatively few polar residues near substrate in the substrate-bound structure. A few additional candidates included E368 and R370. Mutations at these positions retaining steric bulk but removing the polar contact were generated and enzymes incubated with CBGA, however, all mutants retained full conversion to CBCA in endpoint assays (**Figure S8**). Thus, we hypothesized Tcz9 could perhaps directly oxidize substrate without the need to pre-activate *via* deprotonation, a unique mechanism amongst members of the BBE-like enzyme family that oxidize electron-rich resorcinol/phenol derivatives.

To explore Tcz9’s reactivity potential, we replaced the phenolic protons on CBGA (**1**) with methyl groups to generate the dimethoxylated **3** (**Figure 5A**). We monitored the enzyme activity *via* an assay in which H_2_O_2_,^29^ produced as a byproduct of the catalytic cycle, is coupled to the generation of a dye molecule using horseradish peroxidase (**Figure S9**). With CBGA, both Tcz9 and Clz9 produce hydrogen peroxide indicating enzymatic activity, however, with **3**, only Tcz9 generated detectable quantities of hydrogen peroxide, although at a slower rate than with CBGA (**Figure 5B**). This observation indicates that Tcz9 is capable of oxidizing substrates without pre-activation *via* deprotonation. LC-MS analysis revealed that the major product of the reaction had +16 amu from starting material, consistent with hydroxylation. A minor product had a mass of −2 amu from starting material, consistent with a single two-electron oxidation event (**Figure 5C, S10**). To show whether the added hydroxyl group comes from buffer in the water or O_2_, Tcz9 was incubated with substrate in buffer containing ^18^O-labeled water. A corresponding mass shift of +2 amu was observed in the hydroxylation product mass, indicating that the reactive intermediate is quenched through addition of an external nucleophilic water from the buffer (**Figure 5D**). We hypothesized the flavin cofactor of Tcz9 has a higher redox potential than Clz9, and thus it is able to oxidize these “un-activated” substrates. To test this idea, spectroscopic determination of redox potential was carried out with Tcz9 and Clz9 (**Figure S11**). However, the calculated two-electron potential of Clz9 was +95 mV while Tcz9 was slightly *lower*, at +80 mV, signifying that there must be other biochemical differences underlying the differences in observed reactivity.

**Figure 5.**
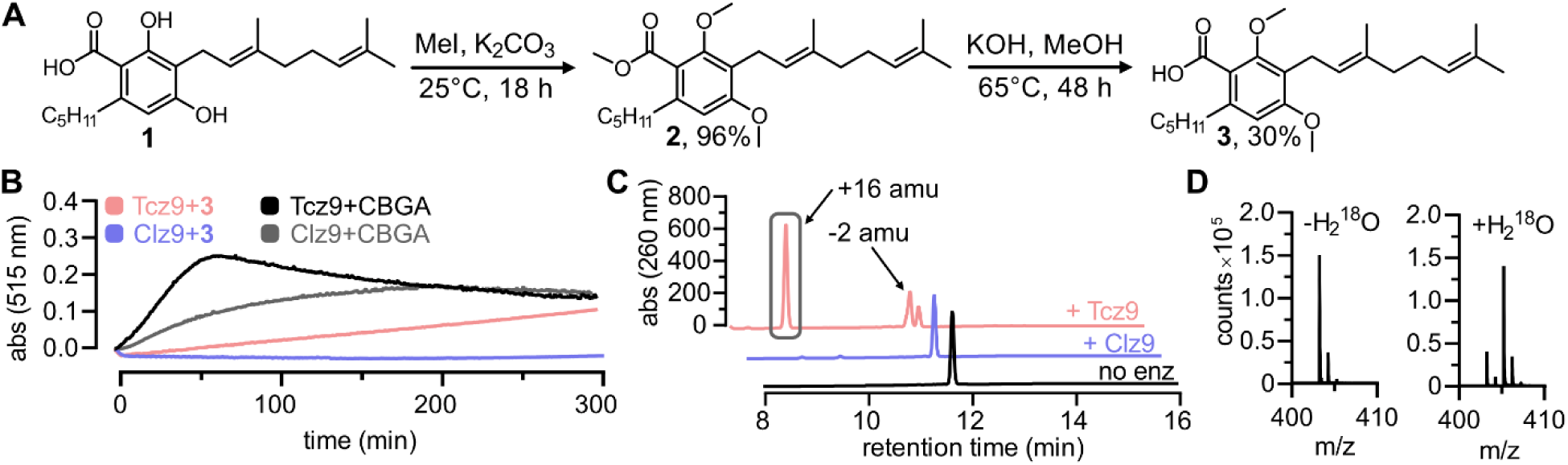
Tcz9 is capable of oxidizing without deprotonation. (A) Synthesis of probe molecule **3** with no protons available for abstraction. (B) Assay data for peroxide formation indicating active cofactor turnover coupled to chromophore formation. (C) LC-MS data showing active turnover of **3** to two distinct products. (D) Inclusion of isotopically labeled water shifted the mass of the major product of Tcz9 incubated with **3** by +2 amu, indicating a hydroxylation product generated by addition of water.

Additional spectroscopic work identified several key differences in the reaction of CBGA with Clz9 or Tcz9. Under anaerobic conditions, the reaction of Clz9 was very slow with both **1** and **3** (**Figure S12A-B, Figure S13A-B**). Upon mixing with **1** (400 mM), a significant change to the flavin absorbance spectrum indicated a large degree of saturation by the substrate. This was followed by a very slow single phase of flavin reduction (k_obs_ = 2.4 h^−1^). Notably, the complex with **1** did not have charge-transfer absorbance indicating that the phenolate of the substrate was not in contact with the isoalloxazine. In contrast to what was observed from the hydrogen peroxide assay, anaerobic incubation of **3** (400 mM) with Clz9 did cause a smaller but still discernable change to the flavin absorbance spectrum indicated the presence of at least some enzyme-substrate complex, again without charge-transfer absorbance. However, the single phase of flavin reduction with **3** was even slower than with **1** (k_obs_ = 0.12 h^−1^). The reactions of Tcz9 with both substrates were notably different. The reaction with **1** was significantly faster than with Clz9, and only the very end of the reaction could be observed in the spectrophotometer (**Figure S12C-D**). From the few data-points that could be collected before reduction was complete, a value of was k_obs_ = ∼30 h^−1^ estimated, but will require rapid-reaction methods for accurate determination. Notably, however, a broad charge-transfer band centered at 600 nm was observed, indicating the stacking of the phenol/phenolate with the isoalloxazine, consistent with the structure. Reaction of **3** was much slower than **1** and without charge-transfer absorbance (**Figure S13C-D**). Two reaction phases were observed (k_obs1_ = 1.2 h^−1^ & k_obs2_ = 0.25 h^−1^), indicating complexity in the reaction mechanism not observed with Clz9 which our current data cannot yet explain. Turnover of **3** may more stringently require productive orientation of the prenyl chain to facilitate conjugation with both the neighboring alkene *and* the electron-rich aromatic. In the Tcz9-CBGA crystal structure, the substrate is positioned to facilitate such conjugation, however, installation of the methoxy substituents could change binding orientation. Furthermore, neither CBGA orientations found in the crystal structure are in the proper configuration for CBCA formation, with both requiring substantial movement of the prenyl chain for productive catalysis (**Figure S5**). Given the rates of Clz9 and Tcz9 with **1**, it is likely that the active substrate conformation is immediately processed and thus, not able to be statically captured using X-ray crystallography.

### Structure-guided enantioselective enhancement of CBCA formation

Despite the reactivity differences between them, both Clz9 and Tcz9 produce a nearly racemic mixture of CBCA enantiomers when incubated with CBGA (**Figure 6**).^18^ In looking at the both apo and substrate-bound structures, this is not necessarily surprising. There is an abundance of space in both active site pockets to accommodate the substrate, as observed in Tcz9 that accommodates two different binding modes (**Figure 3**). The major structural difference between the two binding modes is the orientation of the carboxylic acid in the active site (**Figure S6**). The polar contacts identified between the carboxylate as well as the phenols of the resorcinol provide more information on the relevant interactions between Tcz9 and CBGA, which likely play a role in determining the stereoselectivity of the transformation.

**Figure 6.**
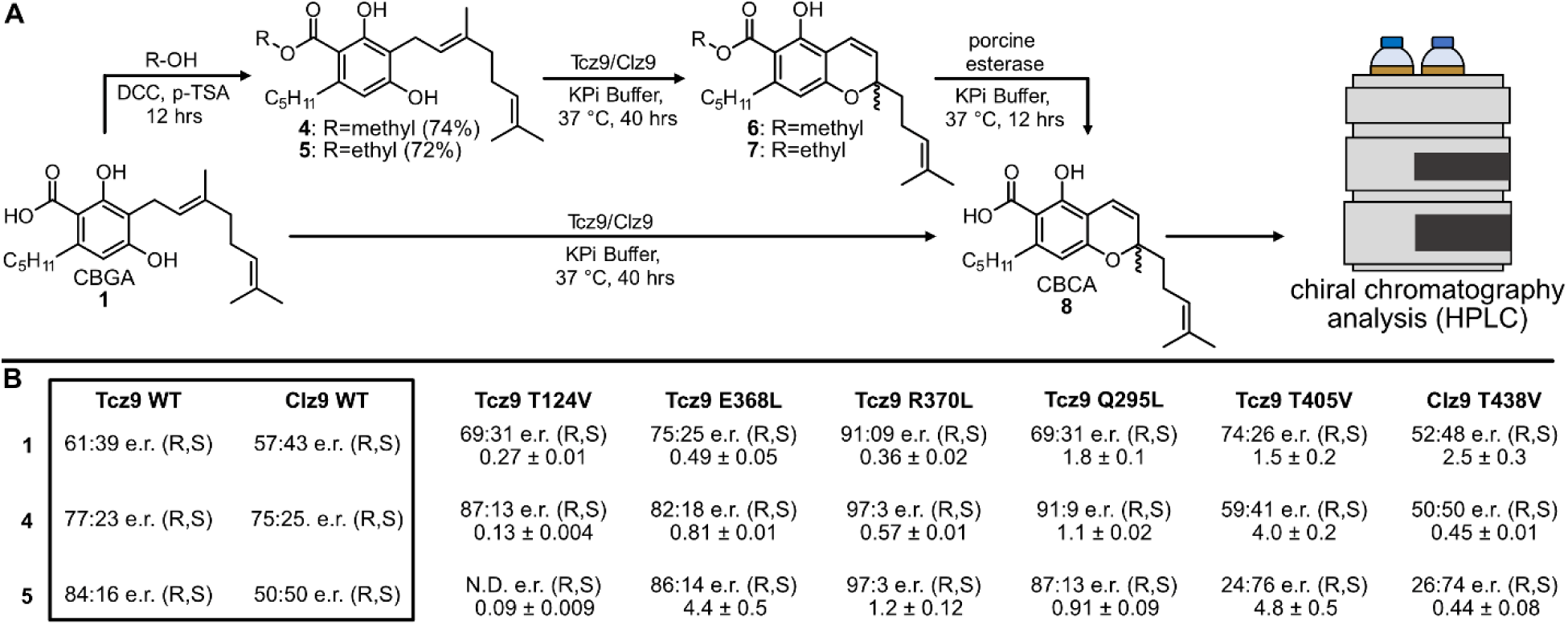
Structure-guided stereochemical biasing of CBCA formation. (A) Methodology for assaying e.r. with a variety of substrates with increasing steric hindrance at the acid position. (B) Tabulated e.r. and relative rate data for all enzymes and substrates tested.

To probe the limits of the active site and the stereoselectivity potential of the reaction, we synthesized and queried a series of CBGA esters against Clz9 and Tcz9 (**Figure 6A**). After enzymatic incubations against methyl, ethyl and isopropyl esters, we removed the ester using porcine liver esterase (PLE) and analyzed the products by LC-MS. As the steric demand increased, the stereoselectivity of the native enzymes improved, peaking at the ethyl ester **5** for Tcz9, which generated the CBCA *R* enantiomer in 5:1 e.r. Clz9, however, had a maximum selectivity of 3:1 e.r. for the *R* enantiomer with the methyl ester **4**, but formed a racemate with the bulkier **5** (**Figure 6B**). In comparison, the stereoselectivity of CBCA synthase from *Cannabis* depends on the strain of plant, but the most selective report indicates that the e.r. in 5:1 with a preference for the *R* enantiomer.^30–34^ The ethyl ester with Tcz9 is capable of matching this selectivity. Increasing the steric bulk beyond ethyl (i.e. isopropyl **s1**) resulted in inefficient to no turnover by Tcz9 and Clz9 (**Figure S14**).

In tandem with modifications to the substrate, we also examined the consequences of removing a polar contact to the substrate or flavin. We thus assayed the mutants that we generated in probing the Tcz9 mechanism (e.g. E368L, R370L, and T405V) as well as two new polar residue mutations (Q295L and T124V) for effects on stereoselectivity of CBCA formation (**Figure 5B, S15**). All mutants displayed higher selectivity for the *R* enantiomer over wild type Tcz9, with the Tcz9-R370L mutant achieving a remarkable 91:9 e.r. This selectivity was higher than what we observed with the most sterically hindered ester (i.e. ethyl ester **5**) and native Tcz9 (**Figure 5B**). When Tcz9-R370L was further incubated with the ester series, this selectivity improved to 97:3 e.r. We similarly observed stereoselectivity improvements, albeit to a lesser degree, across the ester series with most of the other mutants, although in some cases increasing bulk further to **5** resulted in lower (Q295L) to very inefficient (T124V) turnover.

When comparing the relative rates of substrate turnover to wild type Tcz9, the Tcz9-E368L and Tcz9-R370L had lower relative rates (**Figure 5B, S16**). With **5**, however, the relative rate of these two mutants surpassed wild type enzyme. In the structure, the substrate is between the flavin cofactor and these two residues, highlighting their importance in substrate stabilization and positioning in the direction perpendicular to the plane of flavin during catalysis (**Figure 3A**). The Tcz9-T124V mutant exhibited the opposite trend, with decreasing rates as bulk increased, and none of the rates higher than wild type enzyme. In fact, with **5**, the Tcz9-T124V mutant had no appreciable turnover to the corresponding product **7**. In the crystal structure, T124 has a hydrogen bonding interaction with the flavin cofactor, highlighting how positioning of the flavin plays a major role in both selectivity and the rate of catalysis. Tcz9-Q295L had overall higher relative rates when compared to wild type Tcz9, however, the rate decreased as bulk increased. In the structure, Q295 is positioned to the side of where substrate is bound, indicating an importance in determining the positioning in the direction of the plane of the flavin (**Figure 3A**).

The most intriguing trend, however, was observed from the T405V mutant, which flanks residue Q295 in determining positioning of substrate in the plane of the flavin (**Figure 3A**). The initial selectivity with CBGA was improved over wild type Tcz9, and higher than that of the E368, T124, or Q295 mutants. However, when incubated with the ester series, the selectivity decreased for the *R* enantiomer, and ultimately flipped to the *S* enantiomer (3:1 e.r. with **5**). While the *R* enantiomer has been found responsible for the activation of CB2 receptors by CBC, there remains significant unknowns about the biological activity of (*S*)-CBC.^35^ Thus, routes to generate enantiopure material are of high importance. To determine whether this is a unique property of the Tcz9-T405 position, we also tested the stereoselectivity of the corresponding position in Clz9 with a T438V mutant. When incubated with CBGA, **4**, and **5**, we observed the same trend, where incubation of **5** resulted in a 3:1 e.r. for the *S* enantiomer. While the enantioselectivity tells the same story for Tcz9-T405V and Clz9-T438V, the relative rates diverge from each other, with Tcz9-T405V having increasing rates with steric bulk and Clz9-T438V displaying the opposite trend. Even more interestingly, the relative rate of substrate turnover for Clz9-T438V is higher than wild type Clz9 only with CBGA, while Tcz9-T405V displays a higher relative rate than wild type Tcz9 with all three substrates tested. This trend highlights the stark differences in active site architecture between Clz9 and Tcz9, and further indicates that while they may control selectivity in similar manners, there are different dynamics and biochemistry between each of the two with a given substrate.

## Discussion

Clz9 and Tcz9 are amongst a handful of members of the BBE-like enzyme family that perform electrocyclization chemistry to generate the minor cannabinoid CBCA from CBGA.^18^ Our structural data shows that Tcz9 and Clz9 have a high degree of similarity, and thus the differences in biochemistry are notable. The potential capability for Tcz9 to abstract a hydride without the need for substrate “pre-activation” *via* deprotonation has some literature precedent. Phenolic deprotonation is often invoked in *ortho*-quinone methide generation,^7^ however, many BBE-like enzyme are capable of oxidizing substrates that don’t have avenues for a substrate activation step involving deprotonation. For example, the BBE-like enzyme Sol5 from the pathogenic fungus *Alternaria solani* oxidizes a benzylic hydroxyl to an aldehyde without any phenolic protons present on the substrate to deprotonate.^36^ Moreover, the bacterial BBE-like enzyme AknOx from *Streptomyces galilaeus* MA144-M1 also directly oxidizes an alcohol to a ketone.^37^ In this case, the alcohol is on a sugar ring, which is much less electron rich than aromatic species. However, the redox potential and kinetic values suggest that perhaps the major difference between Clz9 and Tcz9 in dictating oxidative chemistry potential is the increased flexibility of the enzyme to move and adapt to substrate.^38^ Both molecular dynamics between the substrate and the enzyme and potentially foundation biochemical differences are likely to play a role in whether given substrates can be oxidized and the rate of oxidation for both Tcz9 and Clz9. Notably, Tcz9 lacks the salt bridge obstructing entrance to the putative oxyanion site and tethering two alpha helices together found in Clz9 (**Figure S2**). Increase enzyme flexibility may also explain why the Tcz9 active site is so resilient to active site changes, as it could potentially move more freely to compensate for unproductive or detrimental interactions. This feature also makes Tcz9 an attractive target for enzyme engineering to expand the chemistry it is capable of performing, and repurpose its cannabinoid cyclase activity toward production of the other two constitutional isomers from the plant (i.e. THC-type and CBD-type cannabinoids).

The growing interest in *Cannabis* and cannabinoids as therapeutics,^39–41^ especially minor cannabinoids like CBCA, necessitates new routes to access enantiopure material. Through both structure and enzyme level engineering, we present a new route to generating enantiopure (*R*)-CBCA, and for generating larger quantities of (*S*)-CBCA (3:1 e.r.), as most tested strains of *Cannabis* preferentially generate the *R* enantiomer.^32, 34^ Our high-resolution substrate-bound structure provided a roadmap for determining the polar contacts between substrate and the active site. We believe that the substrate is likely bound in a local energy minimum that occurs directly before prenyl chain rotation, subsequent to *ortho*-quinone methide formation and electrocyclization. Thermal electrocyclization chemistry proceeds through a disrotatory process,^42^ however, the torquoselectivity of the process is likely dictated by steric occlusion of the prenyl chain upon cyclization.^43^ It is clear from the substrate-bound structure and our spectroscopic interrogation of the Clz9 and Tcz9-mediated CBCA formation chemistry that the dynamics of interactions between the active site and substrate play a considerable role in determining selectivity. We hypothesize that the preferred binding mode leads to *R* enantiomer production, supported by the strong preference for *R* enantiomer formation across mutants and compounds tested. Only in cases where the limits of the active site were pushed through *both* increased bulk *and* active site changes was the *S* enantiomer preferred. With robust rate and biochemical data now in hand, we can begin to probe the dynamics of the process, which is sure to provide key insights that can be applied to better understanding the biochemistry and mechanism of cannabinoid synthases in plants.

Collectively, this work provides new structural data for two intriguing members of the BBE-like enzyme family by interrogating the origins of their cannabinoid cyclization capabilities. In addition, we report here new routes for accessing enantiopure CBCA material, including access to higher quantities of the understudied *S* enantiomer, for biological evaluation. From the biochemistry perspective, the new mechanistic and structural details inform and refine our understanding of the potential of BBE-like enzymes for biocatalytic applications. Further, these insights can be applied to discovery of new bacterial BBE-like enzymes for cannabinoid production and used as inputs for AI models to improve *de novo* design of new cannabinoid cyclases. Lastly, our data informs enzyme engineering efforts toward production of the other two constitutional isomers produced in plants, CBD-type and Δ^9^-THC-type cannabinoids, the subject of ongoing work.

## Methods

### DNA and protein sequences

#### General considerations

DNA sequences were codon-optimized for *Escherichia coli* and ordered from Twist Bioscience and subcloned into a pQE vector using primers and methods described in the **general biological methods** section below.

#### Codon-optimized DNA sequences

##### Clz9

ATGGTTACCGCAGATCCGAGCAGCGAACGTAGCGATATGAATGAAGCAGATGAAGTGAACGAAGTTGATGAACTGAGCGAAACCGGTCAGACCAGCGGCACCAAAGGTAAACGTCCGTTTACAGGTCGTGTTATTGGTCCGGCAGATGGTGAATTTGATGAAGCACGTCGTGTTTGGAATGAATGTTTTGCAGCACGTCCGAAAGAAATTGTTTATTGTGCAGATACCCGTGATGTTGTTCGTGCACTGCGTGAAGTTCGTCAGCGTGGTGGTCCGTTTCGTGTTCGTAGCGGTGGTCATAGCATGAGCGGTCTGAGCGTTCTGGATGATGGTACAGTGCTGGATGTTAGTGGCCTGGATGATATTCAGGTTAGCGAAGATGCAAGCACCGTTACCGTTGGTAGCGGTGCACATCTGGGTGATATTTTTCGTGCCCTGTGGGCACGTGGTGTTACCGTTCCGGCAGGTTTTTGTCCGGAAATTGGTATTGCAGGTCATGTTTTAGGTGGTGGTGCAGGTATTCTGGTGCGTAGCCGTGGTTTTCTGAGCGATCATCTGGTTGCACTGGAAATGGTTGATAGCGAAGGTCGTATTGTTGTTGCAGATCATGATAGTCATCATGAACTGCTGTGGGCAAGCCGTGGTGGTGGCGGTGGTAATTTTGGCATTGCAACCAGCTTTACCCTGCGTACCCAGCCGATTGGTGATGTTACCCTGTTTACCATTGCATGGGATTGGGATCGTGGTGCCGAAGCAATTAAAGCATGGCAAGAATGGCTGGCAACCGCAGATGGTCGCATTAATACACTGTTTATTGCATATCCGCAGGACCAGGATATGTTTGCAGCCCTGGGTTGTTTTGAAGGTGATGCAGCAGAACTGGAACCGCTGATTGCACCGCTGGTTCATGCAGTTGAACCGACCGAAAAAGTTGCAGAAACCATGCCGTGGATTGAAGCACTGAGCTTTGTTGAAACAATGCAGGGTGAAGCCATGAAAGCAACCAGCGTTCGTGCAAAAGGTAATCTGAGTTTTGTTACCGAACCGCTGGGTGATCGTGCCGTTGAAGAAATTAAGAAAGCACTGGCACAGGCACCGAGCCATCGTGCCGAAGTTGTTCTGTATGGTTTAGGTGGCGCAGTTGCAGCAAAAGGTCGTCGTGAAACCGCATTTGTTCATCGTGATGCACCGGTTGCGCTGAATTATCATACCGATTGGGATGATGAAGCCGAAGATGATCTGAATTTTGCCTGGATTCAGAATCTGCGTGCAAGCGTTGCAGCACATACCGAAGGTCGCGGTAGCTATGTTAATACCATTGATCTGACCGTTGAACATTGGCTGTGGGATTATTATGAAGAAAATCTGCCTCGTCTGATGGCCGTTAAAAAGCGTTATGATCCGGAAGATGTTTTTCGTCATCCGCAGAGCATTCCGGTTAGCCTGACCGAAGCAGAAGCAGCCGAACTGGGTATTCCGCCTCATATTGCCGAAGAACTGCGTGCCGCACGTCAGCTGCGT

##### Tcz9

ATGGCAACCCCGAGCGCATTTAGCGGTAGCGTTCTGACACCGGGTGATGATGGTTTTGAAGCAGCACAGGTTACCTGGAATGCATGTTATAGCAGCCGTCCGCGTGAAGTTATGGTTTGTCATGATGCAGCAAGCGTTGCAGAAGCAGTTCGTAGCGTTCGTGAACGTGGTCTGCCGTTTCGTGTGCGTAGCGGTGGTCATAGCATGTGTGGTCTGAGCAATCTGGATGATGGTGTGATTATTGATTTAGGTGGTTTAGGCGGTGTTGAACTGACTCCGGATCGTCAGACCGTTCGTATTGGTGGTGGTGCACGTCTGGCAGATGTGTATAATACCCTGTGGGATCATCGTCTGACAGTTCCGGCAGGCACCTGTCCGCGTATTGGCGTTGGTGGTCATGTGTTAGGTGGTGGTATGGGTGTTCTGAGCCGTAGCCGTGGTGCACTGGTTGATCATCTGACCGCACTGGAAATGGTTGATGCC GAAGGTCGTCTGCTGCGTGTTAGCGAAGATGAAAATCCGGACCTGTTTTGGGCATGTCGTGGTGGTGGCGGTGGTAATTTTGGTATTGTTACCGCATATGAACTGCGTCCGACACCGATTGATGATGTTACCATTTTTACCGTTAGCTGGACCTGGTCACAGCTGCCGGATGCAGTTCGTGCATGGCAGCGTTGGCTGGGTAGCGCAGAAAGCCGTATTAATAGTTTTCTGAGCCTGTTTCCGCAGCAGCAGGATATGGTTGTTGCATTTGGTGTTTTTGATGGTCCGGCAGCAGATTTTCGTCCGCTGCTGGCACCGCTGACCGCAGAAGTTGCACCGGAAGCCGAAGTTGTTGAAGAAGTTCCGTTTATTCAGGCAGTTGATACCGTTGAAGCACTGCAGGGTGAAGCAGCCGCAGCAGAACAGGTTCGTGCACAGGGTAGCAGCGCAATTATTGCAAATCCGCTGAATGATGAAGCCCTGGCAACCCTGCAAGAATTTCTGACCGATCCGCCTAGCCATCGTGCGGAAGTTGCAGTTTATGGCATGGGTGGTGTTATTGGTGAACGTGAACGCGGTGATACCGCATTTGTTCATCGTACCGGTCTGATGGCATTTGAATATCGTACCGATTGGGATAGTCCGGAAGATGATCGTCTGAATCTGGATTGGGTTACCCGTCTGCGTCATGCAATGGCAGAACACACCACCGGTGCAGCATATGTTAATACCATTGATCTGGCCCTGGAAAATTGGCTGTGGGCATATTATGAAGAAAACCTGCCTCGCCTGATGGCAGTTAAACGTCGTTATGATCCGGAAAACGTGTTTCATCATCCGCATAGCATTCCGGGTAGCCTGACAGCCGAAGCAGCCCGTGCACATGGTGTTCCGGAAGCAACCCTGAAACGTCTGCATGATGATGGCCTGCTGGATGGTCCGCTGGAT

#### Protein sequences and Accession Numbers (UniProt)

##### Clz9 (U6A1G7)

MVTADPSSERSDMNEADEVNEVDELSETGQTSGTKGKRPFTGRVIGPADGEFDEARRVWNECFAARPKEIVYCADTRDVVRALREVRQRGGPFRVRSGGHSMSGLSVLDDGTVLDVSGLDDIQVSEDASTVTVGSGAHLGDIFRALWARGVTVPAGFCPEIGIAGHVLGGGAGILVRSRGFLSDHLVALEMVDSEGRIVVADHDSHHELLWASRGGGGGNFGIATSFTLRTQPIGDVTLFTIAWDWDRGAEAIKAWQEWLATADGRINTLFIAYPQDQDMFAALGCFEGDAAELEPLIAPLVHAVEPTEKVAETMPWIEALSFVETMQGEAMKATSVRAKGNLSFVTEPLGDRAVEEIKKALAQAPSHRAEVVLYGLGGAVAAKGRRETAFVHRDAPVALNYHTDWDDEAEDDLNFAWIQNLRASVAAHTEGRGSYVNTIDLTVEHWLWDYYEENLPRLMAVKKRYDPEDVFRHPQSIPVSLTEAEAAELGIPPHIAEELRAAR QLR

##### Tcz9 (A0A7V8NJY5)

MATPSAFSGSVLTPGDDGFEAAQVTWNACYSSRPREVMVCHDAASVAEAVRSVRERGLPFRVRSGGHSMCGLSNLDDGVIIDLGGLGGVELTPDRQTVRIGGGARLADVYNTLWDHRLTVPAGTCPRIGVGGHVLGGGMGVLSRSRGALVDHLTALEMVDAEGRLLRVSEDENPDLFWACRGGGGGNFGIVTAYELRPTPIDDVTIFTVSWTWSQLPDAVRAWQRWLGSAESRINSFLSLFPQQQDMVVAFGVFDGPAADFRPLLAPLTAEVAPEAEVVEEVPFIQAVDTVEALQGEAAAAEQVRAQGSSAIIANPLNDEALATLQEFLTDPPSHRAEVAVYGMGGVIGERERGDTAFVHRTGLMAFEYRTDWDSPEDDRLNLDWVTRLRHAMAEHTTGAAYVNTIDLALENWLWAYYEENLPRLMAVKRRYDPENVFHHPHSIPGSLTAEAARAHGVPEATLKRLHDDGLLDGPLD

### General Biological Methods

#### General considerations

All water used for preparation of bacteria growth media and protein purification buffers was purified from a Milli-Q purification system. All growth media was autoclaved for at least 45 minutes before use. All buffers were vacuum filtered with 0.2 µm nylon filters (Millipore Sigma). All centrifugation steps for protein purification were performed using Beckman Coulter Avanti JXN-26 centrifuges using either the J-8.1 or J18 rotors. Cells were lysed by sonication with a QSonica Q500 ultrasonic processor. Protein samples were analyzed with Tris SDS-PAGE gels (BioRad) and Quick Coomassie stain (Anatrace). All purification steps were performed at room temperature unless otherwise indicated. All Clz9 and Tcz9 enzymes (including mutant enzymes) for mass spectrometry activity assays were purified from plasmids generated as previously described. Clz9 and Tcz9 enzymes for crystallographic analysis had a TEV protease cut site inserted between the His tag and the enzyme, as the His tag was cleaved before crystallization. Plasmids containing the TEV protease cute site were prepared as described below.

#### General cloning methods

Polymerase chain reactions (PCR) methods were performed to amplify the Clz9 and Tcz9 genes to include a TEV protease cut site between the His tag and the enzyme for crystallographic purposes. Amplification was performed using the following primers (three primers total per reaction):

5’-GAGGAGAAATTAACTATGAGAGGATCGCATCACCATCACCATCACgaaaa-3’,
5’-TCACCATCACCATCACgaaaatctatacttccaaggatccATGGTTACCGCAGATCCGAG-3’ and
5’-TGCCGCACGTCAGCTGCGTtaaTCTAGAggcatcaaataaaacgaaaggctcag-3’ for Clz9

And

5’-GAGGAGAAATTAACTATGAGAGGATCGCATCACCATCACCATCACgaaaa-3’,
5’-CATCACCATCACCATCACgaaaatctatacttccaaggatccATGGCAACCCCGAGCGCA-3’ and
5’-CTGCTGGATGGTCCGCTGGATtaaTCTAGAggcatcaaataaaacgaaaggctcagtcga-3’ for Tcz9

All PCR reactions were performed in an Eppendorf Mastercycler thermocycler using the following conditions: 1) 95 °C for 5 min, 2) 95 °C for 30 s, 3) 65 °C for 30 s, 4) 72 °C for 1 min, repeat steps 2-4 twenty times, then 72 °C for 5 min, and hold at 12 °C until retrieved from thermocycler. Reactions were analyzed *via* agarose gel electrophoresis and bands corresponding to appropriate DNA size were cut out and incubated at 50 °C overnight with Agarose Dissolving Buffer (Zymo Research). Melted agarose segments were purified the next day using a Zymo Research Gel Purification Kit. Linearized pQE vectors were generated *via* PCR amplification using the following primers:

5’-TAATCTAGAGGCATCAAATAAAACGAAAGGCTCAG-3’ and
5’-GGATCCGTGATGGTGATGGTGATG-3’

The same method as above was used with a modification to the time held at 72 °C in step 4; for vector amplification this time was increased to 5 min. An aliquot (10 μL) was analyzed *via* gel electrophoresis, and if a band appeared at the appropriate size, *Dpn*I (1.0 μL) was added to each 50 μL reaction and incubated at 37 °C for 3 h. Then, Buffer PB (Qiagen, treated as 2x stock) was added to combined reactions in an Eppendorf tube and DNA purified on a silica gel spin column. DNA was washed with 100% EtOH and Plasmid Wash Buffer (Zymo) before elution with Milli-Q H_2_O. DNA concentrations were recorded using a NanoDrop1000.

Linearized vectors were combined with amplified inserts by Gibson assembly (50 °C for 60 min, 10 μL total reaction volume). The entire reaction was directly transformed into TOP10 *E. coli* chemically competent cells. Colonies were screened using colony PCR to check for presence of genes of interest, and those containing diagnostic bands were expanded overnight in 5 mL LB broth supplements with ampicillin (100 μg/mL) and plasmid DNA was extracted from colonies using a Zymo Research Plasmid Mini-prep Kit. Sequencing analysis confirmed successful plasmid generation (Genewiz).

#### Site-directed mutagenesis methods

Tcz9 and Clz9 plasmids *lacking* the TEV protease cut site were freshly miniprepped and used as template for mutagenic PCR reactions. PCR reactions were prepared with 5 ng/µL of template, 0.2 µM forward OR reverse primer (only 1 per reaction, see Table S2), 25 µL MilliQ H_2_O and 25 µL of 2x Q5 Master Mix (New England Biolabs). PCR amplification was performed using a touchdown methods as described here: 1) 95 °C for 5 min, 2) 95 °C for 30 s, 3) 70 °C for 30 s, 4) 72 °C for 30 s, repeat steps 2-4 ten times, decreasing the temperature of step 3 by 1 degree each cycle, then 5) 95 °C for 1 min, 6) 62 °C for 30 s, 7) 72 °C for 7 min, repeat steps 5-7 eighteen times, and then 72 °C for 10 min, and hold at 12 °C until retrieved from thermocycler. After amplification, 1 μL (20 U) DpnI (New England Biolabs) was added to the PCR mixture and incubated at 37 °C for 3 h. Complementary forward and reverse reactions for the same mutant were combined and processed with a PCR clean-up kit (Qiagen). Following purification, 100-200 ng of DNA was transformed into 70 µL of chemically competent TOP10 *E. coli* cells. Three colonies were selected for miniprep and sequenced to confirm successful incorporation of the desired mutation.

#### Protein expression and purification

The pQE-Tcz9/Clz9 plasmids were transformed into chemically competent *E. coli* BL21(DE3) cells. The transformants were plated on agar plates containing ampicillin. Cells were expanded in LB Amp at 37 °C overnight. The overnight culture (25 mL) was used to inoculate 1 L TB precultures incubated at 37 °C, 200 rpm overnight. The precultures were used to inoculate 1 L TB media supplemented with 0.4% glycerol and 100 µM Ampicillin in 2.4 L Fernbach flasks and incubated at 37 °C to mid-log phase (O.D.∼0.8). The culture was then induced with isopropyl βD-1-thiogalactopyranoside (IPTG, 0.5 mM final concentration), and incubated at 30 °C for 16–18 h. The cells were harvested at 4 °C by centrifugation at 3500 rpm for 10 min. Cell pellets were resuspended in 25 mL of lysis buffer (200 mM Tris Base, 500 mM NaCl, 10% glycerol, pH 7.8). The cells were sonicated using a QSonica Q500 ultrasonic processor and centrifuged at 30,000 *x g* for 45 min at 4 °C. Enzymes were purified from clarified supernatants using a 5 mL HisTrap FF column (Cytiva). HisTrap columns were prepared by flowing 10x column volumes of MilliQ H_2_O, followed by 10x column volumes of lysis buffer by a peristaltic pump at a flow rate of 5 mL/ min. Next, clarified lysate (approx. ∼50 mL) was flowed through, followed by 10x column volumes of wash buffer (20 mM Tris Base, 200 mM NaCl, 30 mM imidazole, pH 7.8) at a flow rate of 2.5 mL/min. Lastly, 5x column volumes elution buffer (20 mM Tris Base, 200 mM NaCl, 250 mM imidazole, pH 7.8) was flowed through (2.5 mL/min) and flow through was collected in 2 mL Eppendorf tubes. Fractions were tested with Bradford Assay (BioRad) for presence of protein. Fractions also appeared yellow, which aided in determining which fractions to test for presence of protein. Fractions containing protein were pooled and concentrated to 2.5 mL using 30 kDa Amicon centrifugal molecular weight cutoff filters. Concentrated proteins were desalted using PD-10 desalting columns (Cytiva) equilibrated with storage buffer (100 mM potassium phosphate, 10% glycerol, pH 7.8) following the manufacturer’s directions. Yields were determined by standard Bradford Assay.

#### Spectroscopic determination of reduction potentials

Reduction potentials were determined by the method of Massey.^44, 45^ Enzyme solutions (1.00 mL) containing 1 mM xanthine, 2 μM benzyl viologen, and phenazine methosulfate were made anaerobic on ice in sealed custom-built cuvettes by 12 cycles of replacing the atmosphere with argon using a Schlenk line essentially as described.^46^ 5 μL – 20 μL of appropriate concentrations of milk xanthine oxidase were in the side-arms of the cuvette. Solutions equilibrated to 25.0° and reactions were initiated by mixing the side-arm contents with the enzyme in the sealed cuvettes. Spectra were recorded in Shimadzu UV-2501PC or UV-2550PC instruments from 700 nm to 300 nm at intervals as short as 90 s or as long as 30 min, as appropriate. The extent of reduction of the dye was obtained from the absorbance at 330 nm or 335 nm (isosbestics for Clz9 and Tcz9, respectively) and the extent of reduction of enzymes were obtained from the absorbance at 490 nm, just beyond the absorbance of phenazine methosulfate. Nernst plots gave the difference between the potential of the indicator dye and the enzyme with slopes close to the theoretical value of 1. 0.1 M KPi, ph 7.0 was used for Tcz9, while this buffer was supplemented with glycerol (20% vol:vol) for Clz9 owing to instability of the reduced enzyme.

#### Kinetics of reduction

Enzyme solutions (1.00 mL) were made anaerobic on ice in sealed custom-built cuvettes by 12 cycles of replacing the atmosphere with argon using a Schlenk line essentially as described.^46^ 10 μL – 20 μL of concentrated of substrate solution, dissolved in DMSO, were in a side-arm of the cuvette. Solutions were then equilibrated to 25.0° and reactions were initiated by mixing the side-arm contents with the enzyme in the sealed cuvettes. Spectra were recorded in Shimadzu UV-2501PC or UV-2550PC instruments from 700 nm to 300 nm at intervals as short as 90 s or as long as 30 min, as appropriate. For the reaction of CBGA, enzyme was in 0.1 M KPi, pH 7.0. For the reactions of the markedly less soluble dimethoxy derivative, 20% DMSO was used as a cosolvent and the buffer concentration was 10 mM. Data were analyzed in KaleidaGraph v 5.0. Spectra were corrected for artifactual baseline shifts by either adding/subtracting a constant or, in the case of the large light-scattering caused by the dimethoxy derivative, subtracting a second-degree polynomial fitted at from 530 nm to 700 nm where the flavin did not absorb. The absorbance at ∼450 nm vs. time was extracted to determine the observed rate constant of flavin reduction. Data were fit to one or two exponentials using the Marquardt-Levenberg routine in KaleidaGraph.

### General Crystallographic Methods

#### Protein production, crystallization, and data collection

Both Tcz9 and Clz9 containing the TEV protease cut site between the His tag and the proteins were expressed using a *E. coli* BL21(DE3) strain. Protein expression was done as a 8-liter fermented culture using a Bioflow 415 bioreactor (New Brunswick Scientific), induced via 0.5 mM IPTG induction for 12-16 hrs, harvested and frozen down at −80 C for further use. Cell pellets were resuspended in lysis buffer (20 mM Tris-HCl, pH 8.0, 150 mM NaCl, 20% glycerol, 1 mM benzamidine, and protease inhibitor cocktail containing PMSF, at a 4:1 (v/w) buffer-to-pellet ratio. Cells were lysed using a disruptor (TS-Series; Constant Systems Inc) at 40 kpsi. Cell debris was removed by centrifugation at 38,000xg, followed by ultracentrifugation at 40,000 RPM using a TI-45 rotor. The solubilized protein was purified by sequential affinity chromatography using Ni Superflow (Qiagen) using fast protein liquid chromatography (AKTA, GE Life Sciences). The running buffer (20 mM HEPES pH 8.0, 200 mM NaCl) contained an increasing imidazole concentration from 20 mM to 300 mM. The protein was desalted on a HiPrep Desalting Column (Cytiva) to remove trace imidazole with a running buffer (20 mM HEPES pH 8.0, 100 mM NaCl). Following desalting, the protein is TEV-cleaved by homemade Tobacco etch virus (TEV) protease overnight at 4 °C at 1:10 ratio to remove the N-terminus His-tagged of the protein. Protealytic cleavage releasing the nascent proteins was confirmed with sodium dodecyl-sulfate polyacrylamide gel electrophoresis (SDS-PAGE). Tcz9 protein was concentrated using a 30 kDa Amicon Ultra membrane pore concentrator (Millipore) and Clz9 was concentrated using 50 kDa concentrator and the subsequent sample was further purified using a size exclusion chromatography (SEC) column (Superdex 200 Increase 10/300 GL) with a low salt buffer (20 mM Tris 8.0 and 50 mM NaCl). SDS-PAGE was used to confirm final protein purity, homogeneity, and molecular weight. The final protein concentration was determined using a Pierce BCA Protein Assay Kit (Thermofisher), followed with quantification by SpectraMax iD5 (Molecular Devices) and NanoDrop Spectrophotometer (NanoPhotometer® NP80).

Both Tcz9 and Clz9 were crystallized using the sitting drop vapor diffusion method at 21 °C. For Tcz9, crystals were obtained using protein concentrated between 20-26 mg/mL using a 30 KDa concentrator (Amicon) against a reservoir solution 0.1M HEPES buffer at pH 7.2, 0.1M calcium acetate, with 19% PEG3350 along with 2M NaCl. Two distinct crystal forms formed in the same drop; one a hexagonal and the other orthorhombic. Although less common, the hexagonal crystal form diffracted significantly higher and, therefore, pursued. To obtain crystals of Clz9, protein was concentrated to ∼30 mg/mL using a 50 KDa concentrator (Amicon) and crystallized against reservoir solution 0.1M HEPES buffer pH 7.3, 0.1M calcium acetate, at 15% PEG3350 with additive screening of 2M NaCl. Only one orthorhombic crystal form was observed. In both cases, crystals appeared in 1-2 weeks and continued to grow larger over time.

Crystals of both Tcz9 and Clz9 were cryoprotected with well solution with 30% glycerol added. To obtain Tcz9 with CBGA, we soaked these crystals in cryoprotection adding 20 µM CBGA for 2 hours. Notably, these crystals initially cracked heavily and remarkably reannealed. Crystals were mounted in a nylon loop and flash cooled in liquid nitrogen. The structure of Tcz9 was originally solved using a home x-ray source (D8 VENTURE, Bruker) and data were processed using the PROTEUM5. For better quality diffraction, we collected data for Tcz9, Tcz9-CBGA, and Clz9 at Advanced Light Source (ALS) beamline (8.2.2) and Stanford Synchrotron Radiation Light source (SSRL) beamline (9-2), at 1.56, 1.93 and 1.95 Å resolution respectively. Diffraction data statistics are shown in Table S1.

#### Structure determination and refinement

The initial protein phases for Tcz9 crystal were determined using the molecular replacement (MR) method using Phaser from the Phenix suite^47^ and the program MOLREP.^48^ The starting template model used as a search probe was derived from AlphaFold2.^49, 50^ The polypeptide chains were manually adjusted into electron density using Coot,^50^ refined with *Phenix.refine*^47^, and validated using MolProbity.^51^ The final *R*_work_ and *R*_free_ values for the refined structures for Tcz9-apo, Tcz9-CBGA and Clz9-apo are 19.2/21.51, 20.31/23.10, and 22.96/25.39 respectively. Interestingly, there were some significant differences between the starting AlphaFold2 derived model and the final refined model, which should further the future accuracy of the structural predictive algorithm. For example, approximately 20% of c-terminal region (residues 447-477) of the AlphaFold 2 model was incorrect in addition to registration problems of the b-sheet regions (usually off by 2 residues) as confirmed by Fo-Fc difference density and validated by subsequent re-refinements. We were able to identify 382 ordered waters. There is also a divalent calcium ion Ca^2+^ that is coordinated in a hexagonal coordination sphere formed by monodentate carboxylate groups of Asp42, nitrogen atom of His41, and one water molecule from two Tcz9 molecules related by a crystallographic 2-fold. Calcium was essential for the formation of this crystal form.

The initial phases used to determine the structure of Clz9 were determined by molecular replacement using a modified model of Tcz9 based on protein sequence identify and homology. A probe model of Clz9 was made using Chainsaw (CCP4) from the Tcz9 model and subsequently used as a search template. Subsequent model-building and refinement corrected the positioning of residues (mostly rotamers) and a loop placement near the active site. We have also found two divalent calcium ions Ca^2+^ in Clz9 structure, one calcium ion has heptahedral coordination with Asp 201, Asp 203 and Ser 204 and surrounding water molecules and other one has octahedral coordination formed by bidentate oxygens of both the carboxylate and carbonyl group (C=O) of Asp 246 and surrounding water molecules.

Tcz9 catalyzes the cyclization of the substrate CBGA to CBCA as product. We were, therefore, surprised to resolve CBGA in the enzyme’s active site from data collected from crystals soaked with the substrate, catching the Tcz9 enzyme right before cyclization. The binding of CBGA to Tcz9 revealed that the substrate binds right below the flavin of the FAD, displacing bound waters in the apo structure. The Fo-Fc and 2Fo-Fc difference densities both support two alternative conformations of CBGA that are flipped 180° relative to each other in the active site. There is density to accommodate the carboxylic acid moiety in both substrate placements, which are related by the pseudo 2-fold symmetry of the CBGA molecule itself.

### General analytical methods

#### General considerations

Reversed phase separations and chiral chromatography was carried out using an Agilent Technologies 1260 Infinity series HPLC equipped with a degasser, quaternary pump, autosamplers, diode array, and fraction collector. For semi-preparative separations, a Kinetex 5 µm C18 100 Å, 250 × 10.0 mm column was used. For chiral analytical scale assays, as well as chiral purifications, a Diacel 5 µm ChiralPak® AD-H 250 x 4.6 mm column was used. For analytical scale reactions analyzed using LC-MS, an Agilent Technologies 1260 Infinity series HPLC equipped with a degasser, binary pump, autosampler, and diode array detector coupled to a 6530 Accurate-Mass Q-TOF MS equipped with a Kinetex 5 µm C18 100 Å, 150 x 4.6 mm column was used. Solvents used were HPLC grade or mass spectrometry grade respectively, and for all separations solvent A was water + 0.1% formic acid (FA) and solvent B was acetonitrile + 0.1% FA. Data was analyzed using MassHunter Workstation software version B.05.01 for high resolution LC-MS analysis and OpenLab CDS for single quadrupole LC-MS analysis, and traces were exported to Microsoft Excel and GraphPad Prism 8 for figures. For HPLC data, data was analyzed using OpenLab CDS software and exported to Microsoft Excel and Graphpad Prism 6 for figures. For purifications, fractions were checked by LC-MS and compounds pooled and dried using a Buchi rotary evaporator and a ThermoFisher SPD140DDA SpeedVac™ Vacuum concentrator.

##### High Resolution Analytical LC-MS Method

0.750 mL/min flow rate, equilibration at 50% solvent B for 2 min followed by a linear 8-minute gradient to 100% solvent B. Hold at 100% solvent B until 17 minutes, followed by a linear 1-minute gradient back to 50% solvent B, and held there for 2 minutes to end the method.

##### Preparative HPLC Method

9 mL/min flow rate, equilibration at 50% solvent B for 2 min followed by a linear 5-minute gradient to 80% solvent B. Followed by a linear 5-minute gradient to 90% solvent, and a 3-minute gradient to 100% B until 13 minutes. Hold for 15 minutes at 100% B, followed by a 2-minute post-run to re-equilibrate back to 50% solvent B.

#### Analytical scale enzyme assays for LC-MS analysis

100 µL reactions comprised of 20 µM Tcz9 or Clz9, 1 µL of 20 mM stock substrate (200 µM final concentration, 1% DMSO final concentration) and potassium phosphate buffer (100 mM potassium phosphate, pH 7.5) were incubated at 37 °C for 20-24 h. The reactions were quenched by 100 µL methanol and centrifuged at 16,000 x g for 2 min prior to filtering with 0.2 µm centrifugal filter. 10 µL of each filtered sample was injected and analyzed *via* high resolution LC-MS and analyzed using the methods described above.

#### Hydrogen peroxide detection assays for colorimetric analysis

100 µL analytical enzyme assays were set up in 96-well plates with 20 μM enzyme, 1 μL of 20 mM stock substrate (200 μM final concentration, 1% DMSO final concentration), 1 μL 10 mM stock 4-aminoantipyrine (AAP, 100 μM final concentration), 5 µL 3,5-dichloro-2-hydroxybenzenesulfonic acid (DCHBS, 1 mM final concentration), and 5 µL horseradish peroxidase (50 mg/mL, 2.5 mg/mL final concentration). Reactions were incubated and absorbance recorded at 515 λ every 30 s for 12 h via Tecan Spark Plate reader. Internal temperature was set to 37 °C with continuous orbital shaking at 80 RPM. Relative rates were calculated by plotting the initial slope of the absorption curves up until pigment saturation and comparing to that of the respective wild-type enzyme in Microsoft Excel.

#### Analytical scale enzyme assays for chiral chromatography analysis

100 μL reactions comprising of 40 μM enzyme, 5 μL stock substrate (1 μM final concentration, 5% DMSO final concentration), and potassium phosphate buffer (100 mM potassium phosphate, pH 7.5) were incubated at 37 °C for 72 h. The reaction was quenched by the addition of 100 μL of methanol and dried down in vacuo. The crude was resuspended in 70 μL of potassium phosphate buffer and 30 μL porcine liver esterase (50 mg/mL) and incubated at 37 °C for 12 h. The reaction was dried down in vacuo and resuspended in 50 μL methanol and centrifuged at 16,000 × *g* for 2 min prior to filtering with a 0.2 μm centrifugal filter. 20 µL of the filtered sample was injected onto the HPLC and analyzed using the methods described above.

### General Chemical Methods

#### General synthetic considerations

Unless stated otherwise, reactions were conducted in oven-dried glassware under an atmosphere of argon using anhydrous solvents. All reagents were obtained from MilliporeSigma, Fisher Scientific, or Alfa Aesar and were used without further purification. Flash column chromatography was performed in normal phase using a Teledyne ISCO CombiFlash® Rf+ Lumen™ system. When noted, preparatory reversed phase purification was performed using an Agilent 1260 Infinity II Preparatory HPLC system equipped with a Kinetex 5 μm C18 (250 × 10.0 mm) column. NMR spectra were obtained on a 500 MHz JEOL NMR spectrometer with a 3.0 mm probe. Data for ^1^H NMR spectra are reported as follows: chemical shift (δ ppm), multiplicity, coupling constant (Hz), and integration. Data for ^13^C NMR spectra are reported in terms of chemical shift. NMR data analysis was performed using MestreNova.

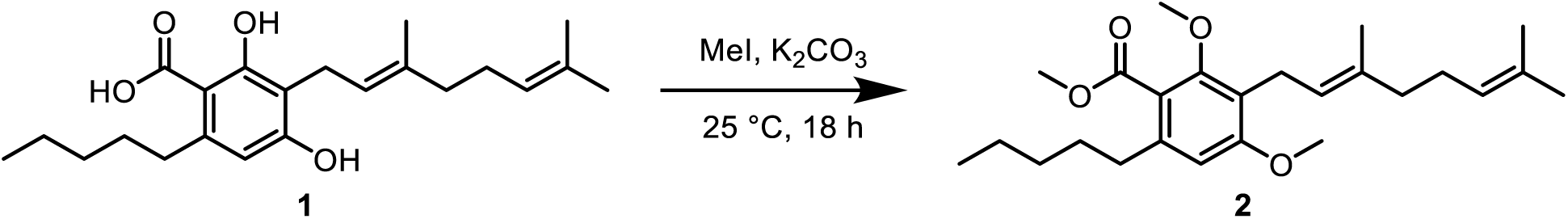

##### Synthesis of 2

To a 50 mL round bottom flask equipped with a magnetic stir bar was added **1** (0.494 g, 1.4 mmol, 1 equiv.), K_2_CO_3_ (0.714 g, 4.9 mmol, 3.5 equiv.) in 12 mL of DMF. The mixture was stirred and MeI (0.30 mL, 4.9 mmol, 3.5 equiv.) was added dropwise to the flask, which was then capped and stirred overnight at room temperature. After 24 h of stirring, the mixture was extracted with EtOAc (2 x 20 mL). The combined organic phases were washed with 4M NaOH in H_2_O (2 x 30 mL) and saturated brine solution (4 x 30 mL). The organic phase was dried over MgSO_4_ and concentrated *in vacuo*. The resulting yellow liquid was dissolved in toluene (2 x 50 mL) and concentrated *in vacuo*. The resulting dark yellow oil was of >98% purity by NMR, and thus was taken on to the next step without further purification (0.515 g, 93% yield).

**^1^H NMR** (500 MHz, CDCl_3_) δ 6.49 (s, 1H), 5.16 (t, *J* = 7.1, 1H), 5.05 (t, *J* = 6.9, 1H), 3.89 (s, 3H), 3.82 (s, 3H), 3.74 (s, 3H), 3.31 (d, *J* = 6.7, 2H), 2.55 (t, *J* = 8.0, 2H), 2.05 (q, *J* = 7.2, 2H), 1.96 (t, *J* = 7.1, 2H), 1.75 (s, 3H), 1.63 (s, 3H), 1.59-1.54 (m, 5H), 1.34-1.29 (m, 4H), 0.88 (q, *J* = 6.8, 5H)

**^13^C NMR** (125 MHz, CDCl_3_) δ 169.3, 159.4, 156.3, 140.1, 135.0, 131.3, 124.4, 122.9, 121.3, 120.8, 107.3, 62.7, 55.7, 52.1, 39.8, 34.0, 31.9, 31.2, 26.7, 25.8, 22.8, 22.5, 22.4, 17.7, 16.2, 14.2, 14.1

**HMRS** (ES−) m/z calc’d for [M – H]^−^: 403.2843, found 403.2867

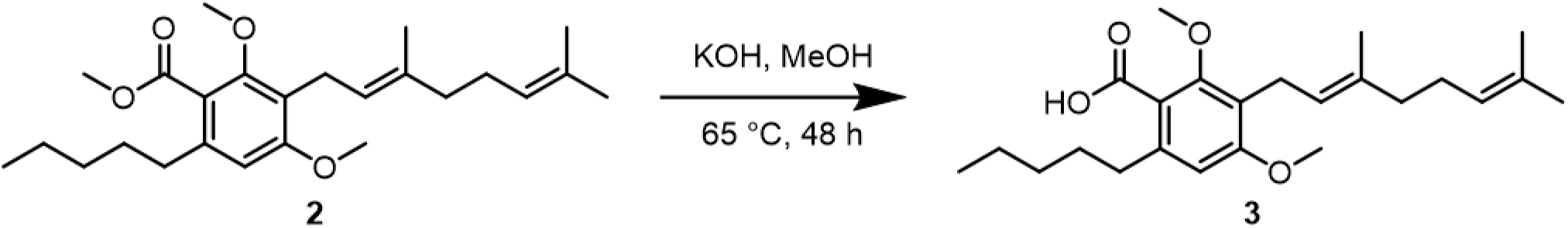

##### Synthesis of 3

To a 50 mL round bottom flask equipped with a magnetic stir bar was added DM-CBGAE (0.102 g, 0.22 mmol, 1 equiv.) as a solution in MeOH (1 mL). A solution of 15% KOH in MeOH (25 mL, 66.7 mmol, 300 equiv.) was then added to the flask and the solution was refluxed at 65 °C for 18 h. The reaction was cooled to room temperature and 10 M HCl was added to reach pH ∼2, and the resulting solution was extracted with CH_2_Cl_2_ (3 x 20 mL). The combined organic phases were washed with H_2_O (1 x 20 mL). The organic phase was dried over MgSO_4_, filtered, and concentrated *in vacuo*. The crude clear oil was dissolved in MeOH, filtered and purified by preparatory reverse phase HPLC using the <u>preparatory HPLC method</u> above. Compound **3** was isolated as a pale purple residue (0.0272 g, 30% yield), and unreacted starting material was also recovered (0.0672 g, 66% recovery) for a total material recovery of 96%.

**^1^H NMR** (500 MHz, CDCl_3_) δ 6.55 (s, 1H), 5.17 (t, *J* = 6.4, 1H), 5.05 (t, *J* = 6.8, 1H), 3.85 (s, 3H), 3.83 (s, 3H), 3.34 (d, *J* = 6.7, 2H), 2.77 (t, *J* = 8.0, 2H), 2.05 (q, *J* = 6.8, 2H), 2.00-1.96 (m, 2H), 1.76 (s, 3H), 1.66-1.60 (m, 5H), 1.57 (s, 3H), 1.38-1.33 (m, 4H), 0.90 (t, *J* = 7.0, 3H)

**^13^C NMR** (125 MHz, CDCl_3_) δ 160.1, 157.2, 135.3, 131.4, 124.4, 122.7, 121.5, 118.2, 108.3, 63.1, 55.8, 39.8, 34.5, 31.9, 31.7, 26.7, 25.8, 22.8, 22.6, 17.7, 16.2, 14.1

**HMRS** (ES−) m/z calc’d for [M – H]^−^: 387.2541, found 387.2548

#### General procedure for esterification

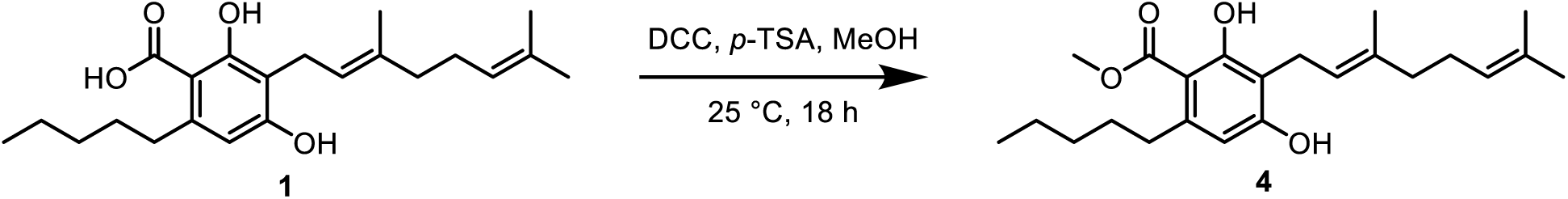

This procedure was adapted from literature precedent.^52^ To a 20 mL scintillation vial equipped with a magnetic stir bar was added **1** (0.090 g, 0.25 mmol, 1 equiv.), dicyclohexylcarbodiimide (0.20 g, 0.97 mmol, 4 equiv.), catalytic *p*-toluensulfonic acid (0.005 g, 0.03 mmol, 0.12 equiv.) and MeOH (5 mL). After 18 h, the solution was worked up by evaporation and redissolved in toluene (5 mL). To precipitate urea the solution was cooled in a freezer (−20 °C) for 2 h. The solution was filtered on a sintered glass filter and concentrated *in vacuo*. The crude was purified by reversed phase preparatory HPLC using the method described above to afford **4** (0.07 g, 74% yield).

**^1^H NMR** (500 MHz, CDCl_3_) δ 6.23 (s, 1H), 5.28 (t, *J* = 5.9, 1H), 5.06 (t, *J* = 6.6, 1H), 3.43 (d, *J* = 7.3, 2H), 2.80 (t, *J* = 7.8, 2H), 2.13-2.03 (m, 4H), 1.81 (s, 3H), 1.59 (s, 3H), 1.55-1.48 (m, 2H), 1.36-1.31 (m, 4H), 0.91 (t, *J* = 6.7, 3H)

**^13^C NMR** (125 MHz, CDCl_3_) δ 172.6, 162.7, 159.6, 145.9, 138.9, 132.1, 124.0, 121.6, 111.7, 111.0, 104.6, 52.0, 39.8, 36.9, 32.2, 31.7, 26.5, 25.8, 22.7, 22.2, 17.8, 16.3, 14.2

**HRMS** (ES−) m/z calc’d for C_23_H_33_O_4_ [M-H]^−^ : 373.2384; found 373.2383

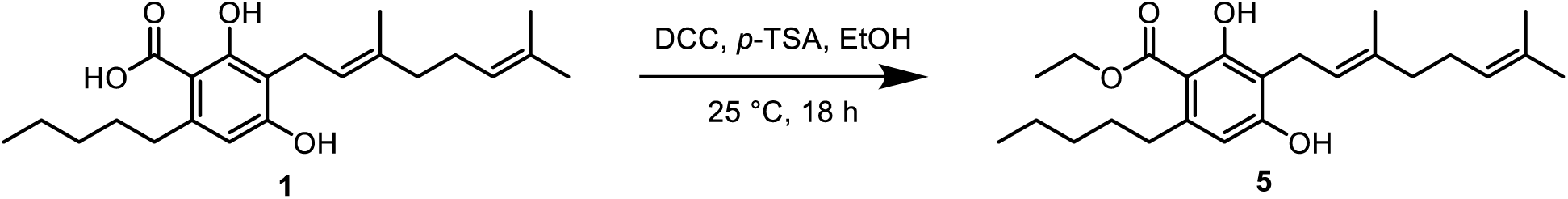

Prepared according to the general procedure for esterification, using **1** (0.095 g, 0.25 mmol, 1 equiv), dicyclohexylcarbodiimide (0. 17 g, 0.82 mmol, 4 equiv), catalytic *p*-toluensulfonic acid (0.005 g, 0.03 mmol, 0.12 equiv), and EtOH (5 mL) to give **5** (0.07 g, 72% yield).

**^1^H NMR** (500 MHz, CDCl_3_) δ 6.23 (s, 1H), 5.28 (t, *J* = 7.2, 1H), 5.06 (t, *J* = 6.7, 1H), 4.40 (q, *J* = 7.1, 2H), 3.43 (d, *J* = 7.2, 2H), 2.82 (t, *J* = 7.8, 2H), 2.14-2.04 (m, 4H), 1.81 (s, 3H), 1.68 (s, 3H), 1.59 (s, 3H), 1.57-1.51 (m, 2H), 1.41 (t, *J* = 7.2, 3H), 1.35-1.32 (m, 4H), 0.90 (t, *J* = 7.2, 3H)

**^13^C NMR** (125 MHz, CDCl_3_) δ 172.2, 162.7, 159.5, 145.9, 138.9, 132.1, 124.0, 121.7, 121.6, 111.7, 110.9, 104.7, 61.4, 39.8, 37.0, 32.2, 31.9, 26.5, 25.8, 22.2, 17.8, 16.3, 14.2

**HRMS** (ES−) m/z calc’d for C_24_H_35_O_4_ [M-H]^−^ : 387.2541; found 387.2548

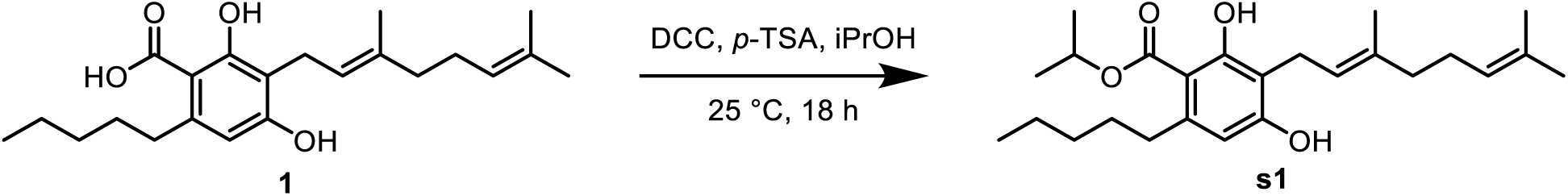

Prepared according to the general procedure for esterification, using **1** (0.095 g, 0.26 mmol, 1 equiv.), dicyclohexylcarbodiimide (0. 18 g, 0.87 mmol, 4 equiv), catalytic *p*-toluensulfonic acid (0.005 g, 0.03 mmol, 0.12 equiv), and isopropanol (5 mL) to give **s1** (0.03 g, 36% yield)

**^1^H NMR** (500 MHz, CDCl_3_) δ 6.22 (s, 1H), 5.30 (q, *J* = 6.3, 1H), 5.27 (t, *J* = 5.9, 1H), 5.05 (t, *J* = 6.9, 1H), 3.43 (d, *J* = 7.2, 2H), 2.82 (t, *J* = 8.0, 2H), 2.12-2.04 (m, 4H), 1.81 (s, 3H), 1.67 (s, 3H), 1.59 (s, 3H), 1.59-1.51 (m, 2H), 1.39 (d, *J* = 6.3, 6H), 1.35-1.32 (m, 4H), 0.91-0.88 (t, *J* = 7.2, 3H)

**^13^C NMR** (125 MHz, CDCl_3_) δ 171.8, 162.7, 159.4, 145.4, 139.1, 132.2, 123.9, 121.6, 111.5, 110.9, 105.1, 69.2, 39.8, 37.1, 32.3, 32.1, 26.5, 25.8, 22.9, 22.2, 22.1, 17.6, 16.4, 14.2

**HRMS** (ES−) m/z calc’d for C_25_H_37_O_4_ [M-H]^−^ : 401.2697 found 401.2704

## Acknowledgments

The authors acknowledge the use and assistance from the Stanford Synchrotron Radiation Laboratory (SSRL) and Advanced Light Source (ALS) for synchrotron data collection and processing. Funding was generously provided by the National Center for Complementary and Integrative Health (1R01-AT012641) to B.S.M. and G.C. and the National Institutes of General Medical Sciences (1F32-GM150232-02) to A.C.L. We thank Trevor Purdy for helpful advice. We thank Amyris for providing CBGA substrates.

## Data and Materials Availability

The X-ray crystallographic coordinates for three structures Tcz9, Tcz9-CBGA and Clz9 have been deposited in the Protein Data Bank (RCSB PDB) under accession codes: 9P1R, 9P1O and 9P1P.

