## Supplemental information for "Structural and biochemical basis for cannabinoid cyclase activity in marine bacterial flavoenzymes"

### TABLE OF CONTENTS

**Table S1-S2** **S2-S3**

**Figures S1-S16** **S4-S19**

**NMR spectra** **S20-S29**

**Table S1.** Clz9, Tcz9 and Tcz9-CBGA data collection and refinement statistics

| **Data Collection** | **TCZ9-APO** | **TCZ9-CBGA** | **CLZ9-APO** |
|---|---|---|---|
| Beamline | ALS (8.2.2) | SSRL 9-2 | SSRL 9-2 |
| Wavelength (Å) | 1.00002 | 0.97946 | 0.97946 |
| Resolution (Å)[a] | 48.57-1.56 (1.59-1.56) | 81.76-1.93 (1.96-1.93) | 85.19-1.95 (1.98-1.95) |
| Space group | P $6_1$ 2 2 | P $6_1$ 2 2 | P $4_1$ $2_1$ 2 |
| Unit cell (Å) | 94.93, 94.93, 240.77 | 94.29, 94.29, 237.06 | 101.39 101.39 157.10 |
| ( °) | 90,90,120 | 90, 90, 120 | 90, 90, 90 |
| Total reflections | 1684297 | 940708 | 810119 |
| Unique reflections | 90005 (37,60) | 47637 (2354) | 60140 (2924) |
| Multiplicity | 18.7(12.7) | 19.7 (15.2) | 13.5 (11.5) |
| Completeness (%) | 98.2 (85.7) | 100.0(100.0) | 100.0(99.6) |
| Mean (I)/ $\sigma_I$ | 26.4 (0.9) | 19.4 (0.5) | 18.0 (0.7) |
| Rmerge [b](%) | 6.8 (>100) | 7.4 (>100) | 9.0 (>100) |
| $R_{meas}$[c](%) | 7.2(>100) | 7.6 (>100) | 9.4 (>100) |
| $R_{pim}$[d](%) | 2.2(>100) | 1.7 (>100) | 2.5 (>100) |
| $CC_{1/2}$[e](%) | 100.00 (30.4) | 99.9 (33.2) | 100.00 (34.9) |
| **Refinement Statistics** | | | |
| Refinement resolution (Å)[a] | 48.57-1.56 (1.61 – 1.56) | 77.21 - 1.93 (2.00 - 1.93) | 50.69 - 1.95 (2.02 - 1.95) |
| # reflections in refinement (work/free) | 88,805/ (7,817) | 47,217/ (4,325) | 59,974/ (5,738) |
| $R_{work}$ (%) | 19.20 | 20.31 | 22.96 |
| $R_{free}$ (%) | 21.51 | 23.10 | 25.39 |
| # Protein atoms | 3,610 | 3,631 | 3,599 |
| # Waters | 382 | 107 | 210 |
| # Protein residues | 472 | 472 | 468 |
| Bond r.m.s. deviation (Å) | 0.015 | 0.006 | 0.003 |
| Angle r.m.s. deviation (°) | 1.87 | 0.94 | 1.25 |
| Ramachandran favoured, allowed, outliers (%)[1] | 97.45, 2.34, 0.21 | 96.81, 2.77, 0.43 | 95.92, 4.08, 0.00 |
| Wilson B (Å$^2$) | 26.36 | 49.26 | 41.27 |
| Average B (Å$^2$) | 41.88 | 70.05 | 57.62 |
| Ligand c[g] | 31.33 | 76.13 | 45.04 |
| Protein | 41.36 | 70.01 | 57.83 |
| PDB ID | 9P1R | 9P1O | 9P1P |

[a]Numbers in parentheses are for highest resolution shell

[b]$R_{merge} = \Sigma_{hkl} \Sigma_{i=1,n}|I_i(hkl) - \langle I(hkl)\rangle| / \Sigma_{hkl} \Sigma_{i=1,n} I_i(hkl)$

[c]$R_{meas} = \Sigma_{hkl} \sqrt{(n/n-1)}\, \Sigma_{i=1,n}|I_i(hkl) - \langle I(hkl)\rangle| / \Sigma_{hkl} \Sigma_{i=1,n} I_i(hkl)$

[d]$R_{pim} = \Sigma_{hkl} \sqrt{(1/n-1)}\, \Sigma_{i=1,n}|I_i(hkl) - \langle I(hkl)\rangle| / \Sigma_{hkl} \Sigma_{i=1,n} I_i(hkl)$

[e]$CC_{1/2}$ = Pearson Correlation Coefficient between two random half datasets

[f]Number of unfavorable all-atom steric overlaps ≥ 0.4Å per 1000 atoms

[g] Bound ligand c are FAD, CBGA and Ca2+

**Table S2.** Primers for site-directed mutagenesis of Clz9 and Tcz9.

| number | direction | mutant | primer |
|---|---|---|---|
| 1 | forward | Clz9-Y435F | CATACCGAAGGTCGCGGTAGCtttGTTAATACCATTGATC |
| 2 | reverse | Clz9-Y435F | GATCAATGGTATTAACaaaGCTACCGCGACCTTCGGTATG |
| 3 | forward | Tcz9-Y402F | ACACACCACCGGTGCAGCAtttGTTAATACCATTG |
| 4 | reverse | Tcz9-Y402F | CAATGGTATTAACaaaTGCTGCACCGGTGGTGTGT |
| 5 | forward | Clz9-Y374F | TGCCGAAGTTGTTCTGtttGGTTTAGGTGGCGCAGTT |
| 6 | reverse | Clz9-Y374F | AACTGCGCCACCTAAACCaaaCAGAACAACTTCGGCA |
| 7 | forward | Tcz9-Y342F | GCGGAAGTTGCAGTTtttGGCATGGGTGGTGTT |
| 8 | reverse | Tcz9-Y342F | AACACCACCCATGCCaaaAACTGCAACTTCCGC |
| 9 | forward | Clz9-T438V | CGCGGTAGCTATGTTAATgttATTGATCTGACCGTTGAACATTG |
| 10 | reverse | Clz9-T438V | CAATGTTCAACGGTCAGATCAATaacATTAACATAGCTACCGCG |
| 11 | forward | Tcz9-T405V | ACCGGTGCAGCATATGTTAATgttATTGATCTGGCCCTGG |
| 12 | reverse | Tcz9-T405V | CCAGGGCCAGATCAATaacATTAACATATGCTGCACCGGT |
| 13 | forward | Tcz9-Q295L | TGATACCGTTGAAGCACTGttaGGTGAAGCAGCCGCAGCAGAACAGGTTC |
| 14 | reverse | Tcz9-Q295L | GAACCTGTTCTGCTGCGGCTGCTTCACCtaaCAGTGCTTCAACGGTATCA |
| 15 | forward | Tcz9-R370L | ATGGCATTTGAATATctgACCGATTGGGATAGTCC |
| 16 | reverse | Tcz9-R370L | GGACTATCCCAATCGGTcagATATTCAAATGCCAT |
| 17 | forward | Tcz9-E368L | GGTCTGATGGCATTTctgTATCGTACCGATTGG |
| 18 | reverse | Tcz9-E368L | CCAATCGGTACGATAcagAAATGCCATCAGACC |
| 19 | forward | Tcz9-T124V | CAGTTCCGGCAGGCgttTGTCCGCGTATTGGC |
| 20 | reverse | Tcz9-T124V | GCCAATACGCGGACAaacGCCTGCCGGAACTG |

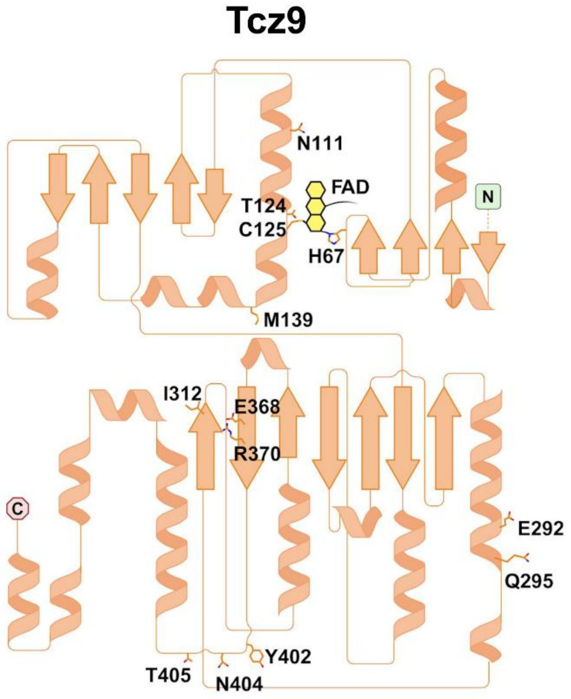


**Figure S1. 2D topology diagram of secondary structure.** Topology for Tcz9 (pink, left) and Clz9 (purple, right). FAD is indicated along with key residues in the active site. Both enzymes are topologically nearly identical. Image was made using Biorender.

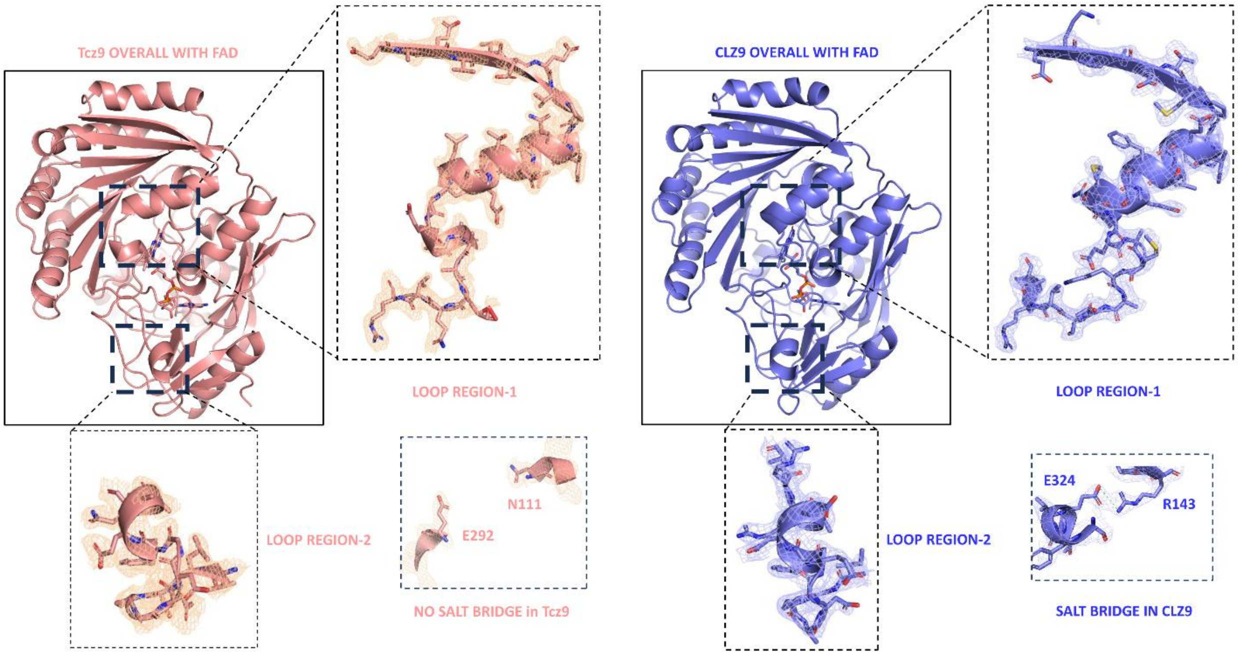


**Figure S2. Electron density for Tcz9 and Clz9.** Shown are the loop regions 1 and 2 with superimposed 2Fo-Fc electron densities (1 sigma) for Tcz9 (left) and Clz9 (right). The electron density for residues that form the salt bridge for Clz9 (Residues E324 and R143) are shown along with corresponding residues in Tcz9 that do not form salt bridge (E292 and N111).

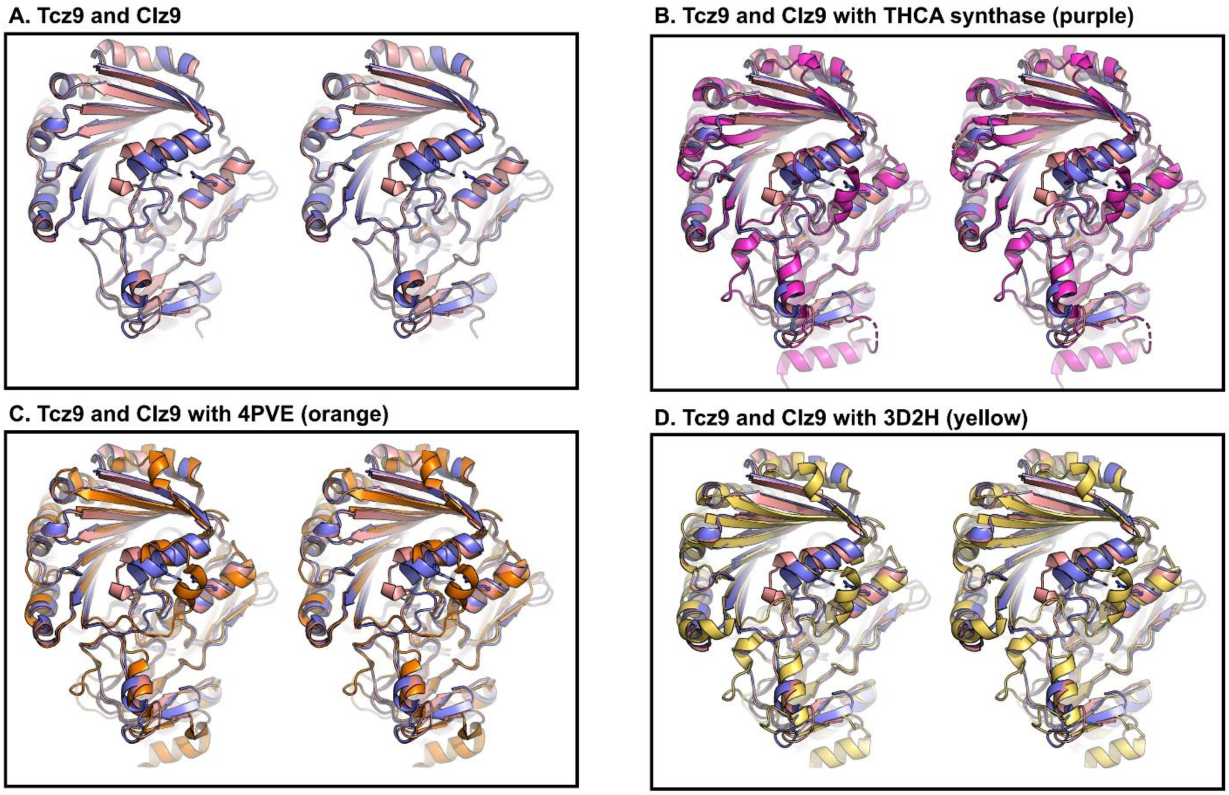


**Figure S3. Structural comparisons with BBE-like enzymes.** (A) Comparison of Tcz9 and Clz9 as reference (same view as in Fig 2A). Structural comparison of Tcz9 and Clz9 with (B) THCA (violet), (C) Phl p4 (4PVE) (orange), and (D) BBE-enzyme from *E. californica* (3D2H) (yellow).

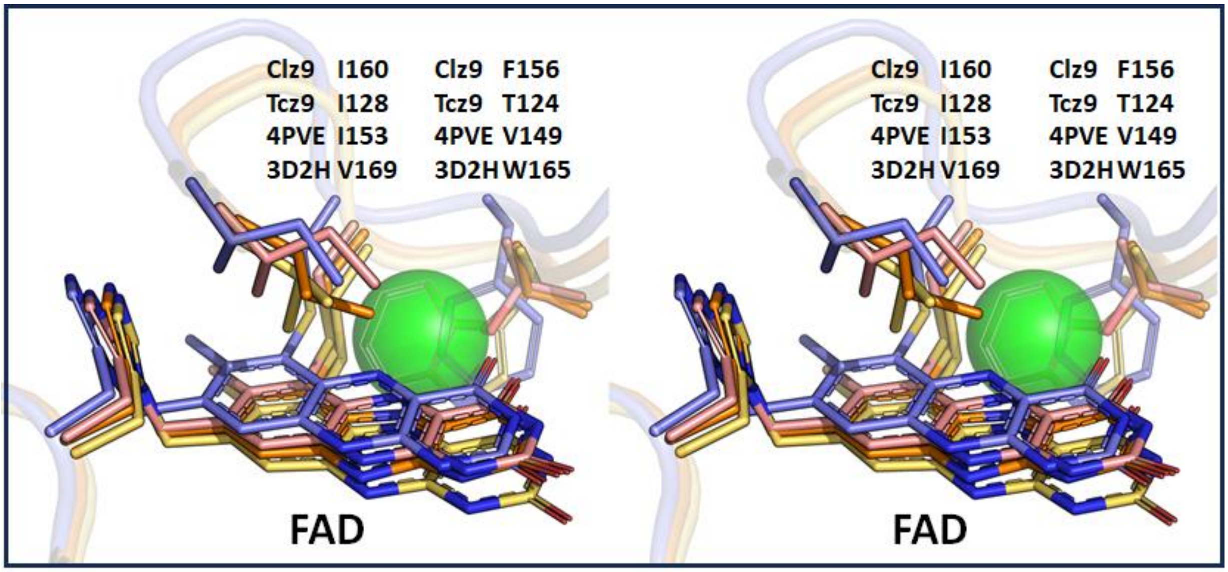


**Figure S4. Putative "Oxyanion hole".** Stereo-view of the putative "oxyanion hole" location is indicated as a transparent green sphere, superimposed with the models of Phl P4 (yellow), the BBE-enzyme from *E. californica* (orange), Tcz9 (pink), and Clz9 (purple). The putative "gate-keeper" residue position identified to play a role in Phl p4 (4PVE-I153), and the BBE-enzyme from *E. californica* (3D2H-V169) are shown along with the corresponding residues in Tcz9-I128 and Clz9-I160. The isoleucine positions in Tcz9 and Clz9 are displaced from the oxyanion hole. The relative positioning of flavins are indicated.

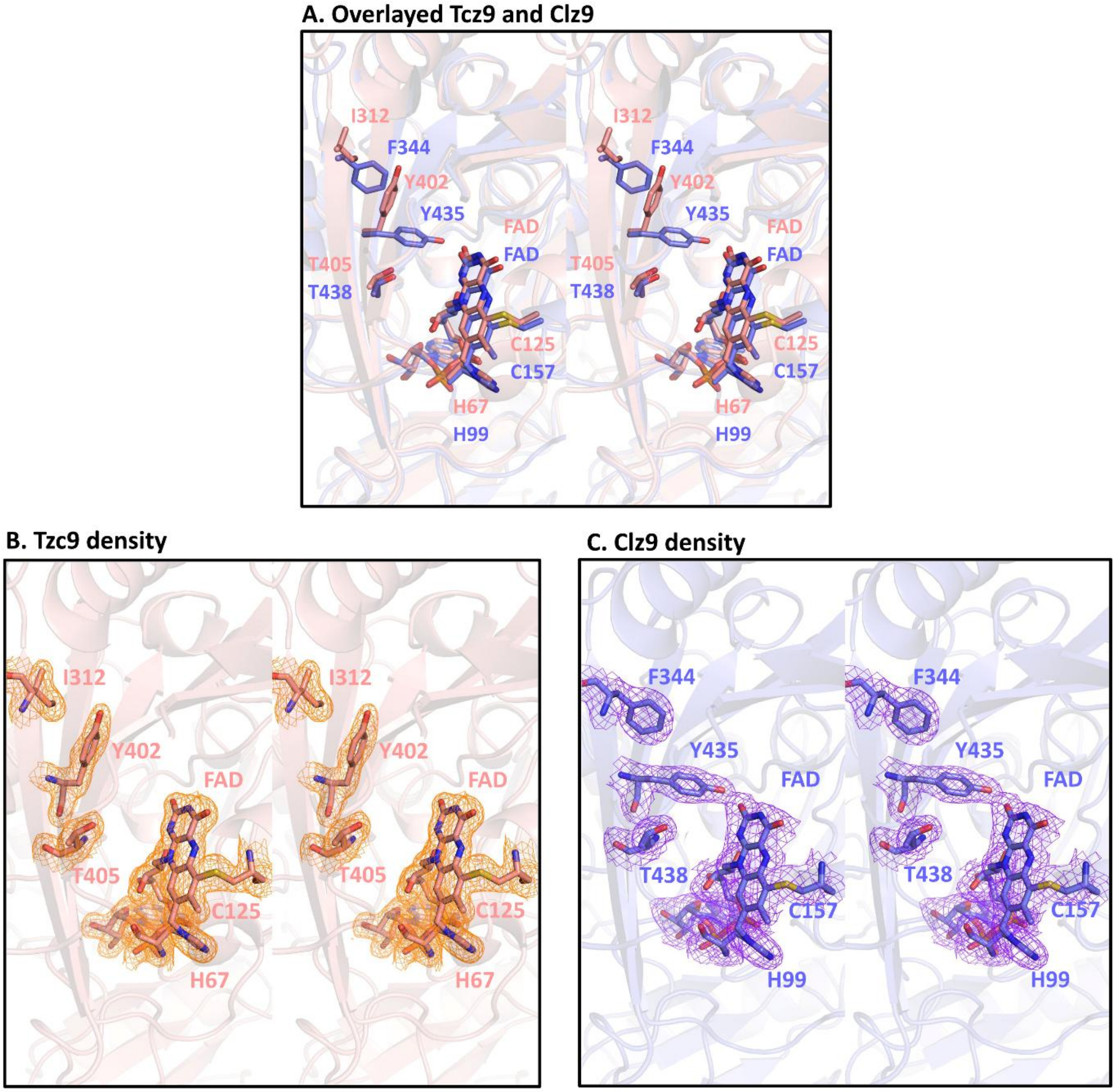


**Figure S5. Stereo close-up views of Y/FxN motif of Tcz9 (pink) and Clz9 (purple).** (A) Shown are the relative positional differences in the placement of Tcz9Y402 and Clz9-Y435 caused by Tcz9-I312 and Clz9-F344. (B-C) Corresponding stereo-views of the 2Fo-Fc density (contoured at 1$\sigma$) for Tcz9 and Clz9, respectively.

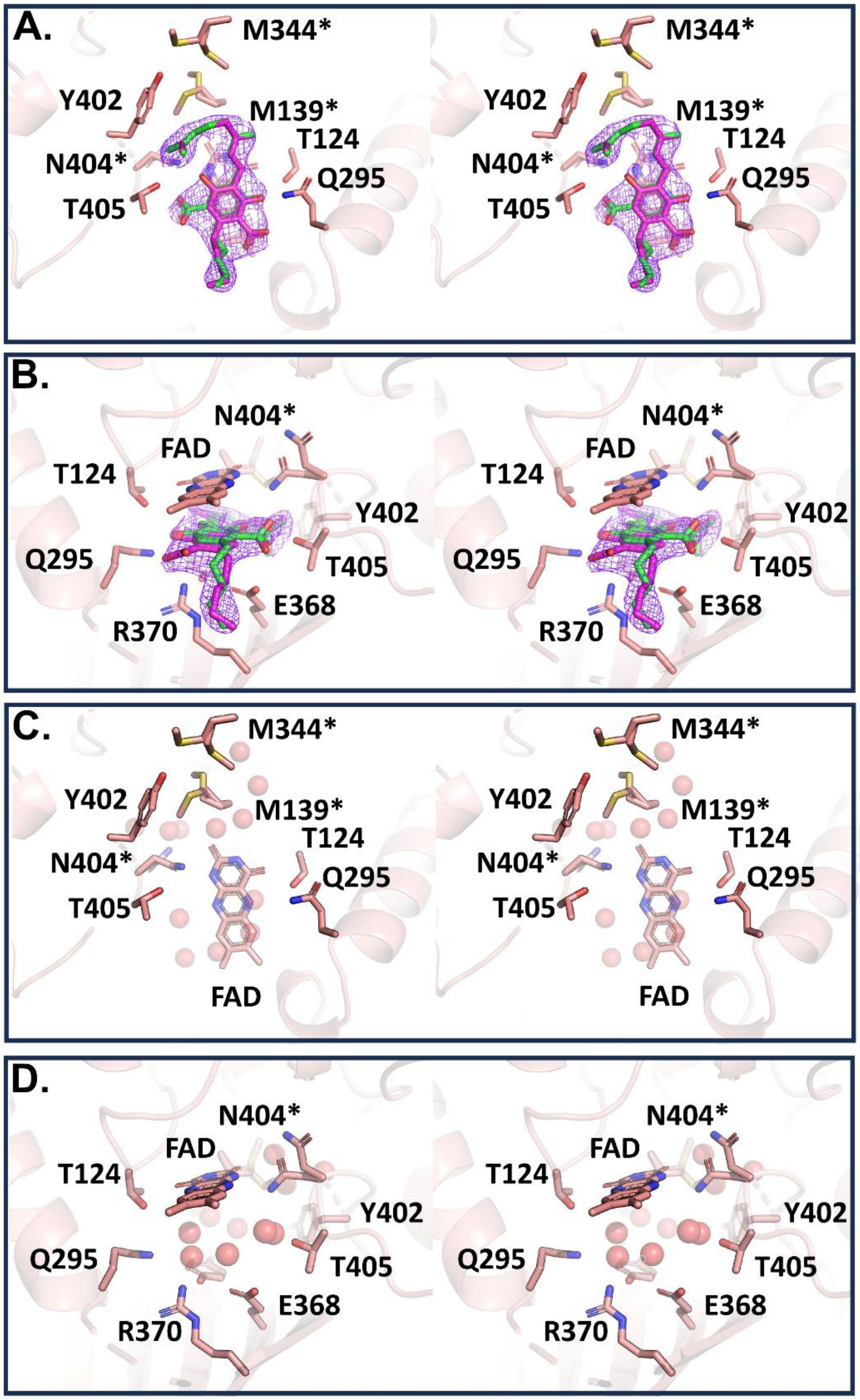


**Figure S6. CBGA displaces bound water in Tcz9.** Stereo views are shown of Tcz9 with CBGA bound and apo Tcz9. Views are shown (A-B) perpendicular to and (C-D) nearly parallel with the plane of the flavin. 2Fo-Fc electron density surrounding both alternative conformations of CBGA used in crystallographic refinement are contoured at 0.6σ. Bound ordered waters (B-D; red spheres) in the apo Tcz9 structure that are displaced by CBGA binding shown in the same relative views as in A and C. Residues denoted with an asterisk have alternative conformations in the electron density map.

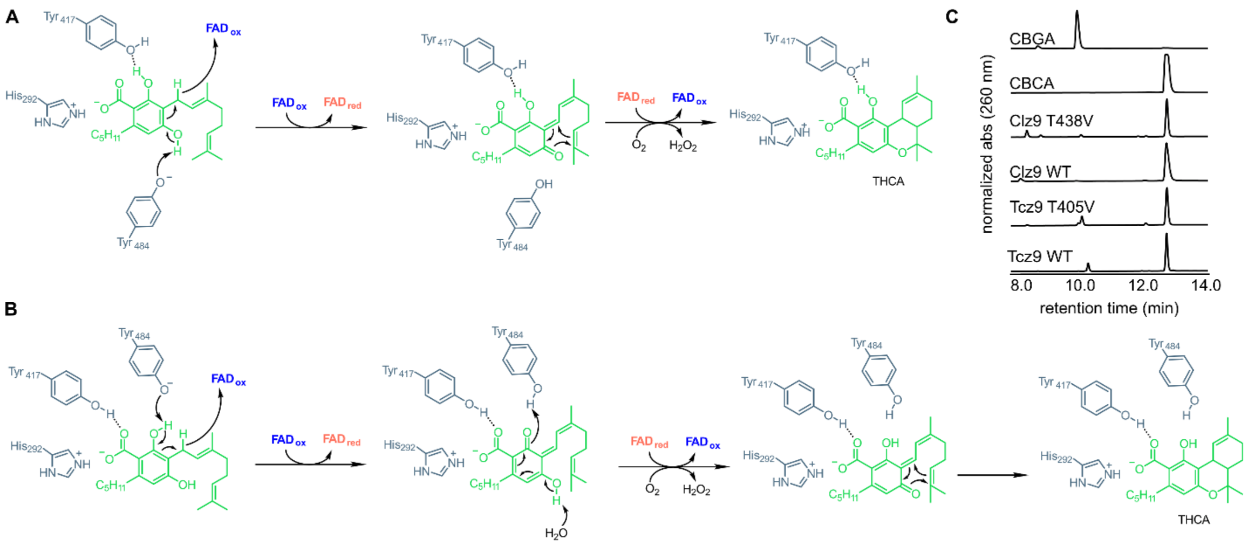


**Figure S7.** Proposed mechanisms for cyclization of cannabinoids in plants. (A) Initial proposed mechanism based on mutagenesis work from Shoyama *et. al.*[8] (B) Competing mechanism published by Villard *et al.*[28] (C) Threonine homologous to catalytic tyrosine in plant enzymes is not necessary for catalysis by Clz9 and Tcz9.

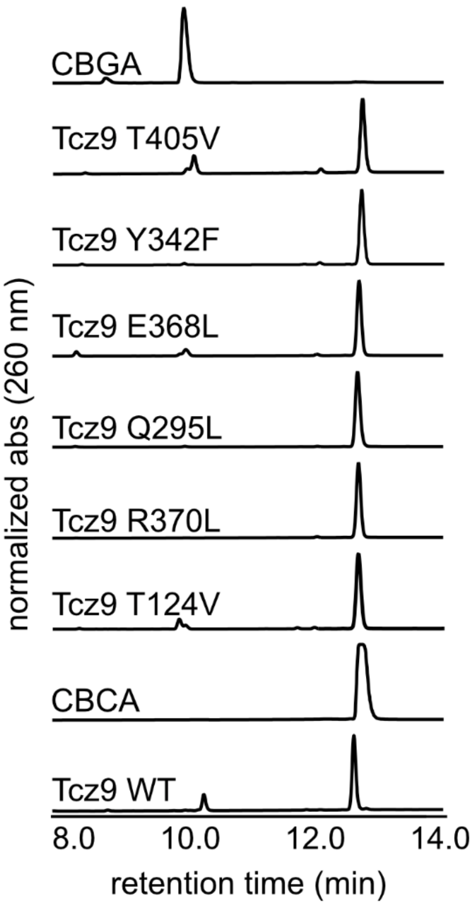


**Figure S8.** Polar residues in the active site of Tcz9 are not required for substrate activation and catalysis in CBCA formation.

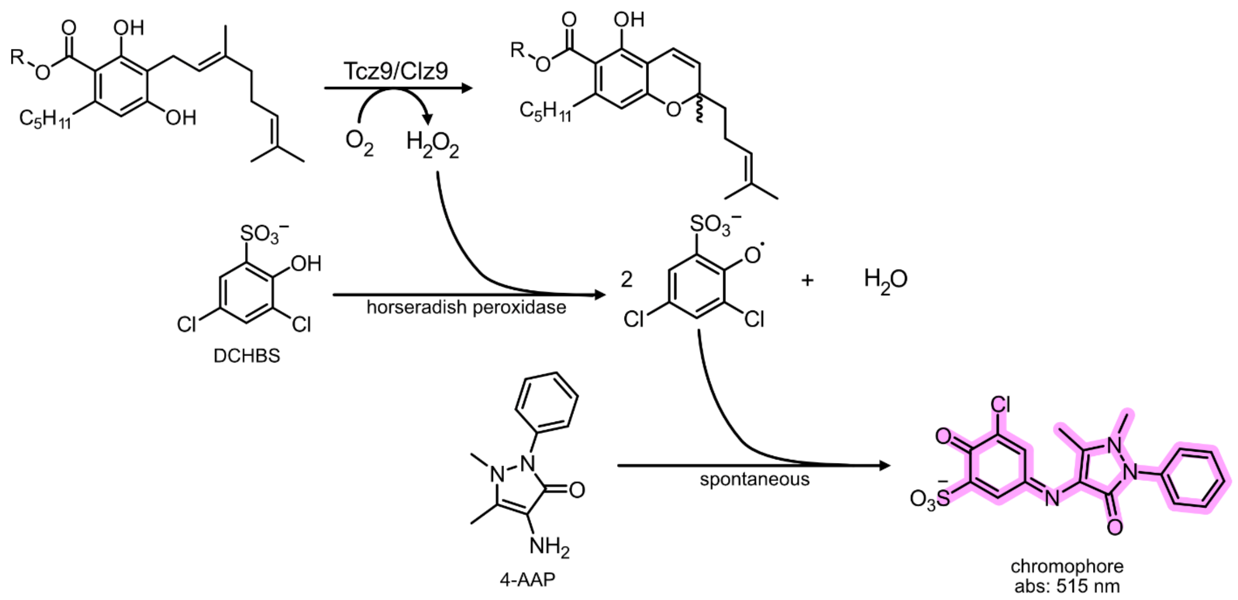


**Figure S9.** Production of hydrogen peroxide during CBCA formation can be coupled to production of an absorbent dye through horseradish peroxidase-mediated turnover.

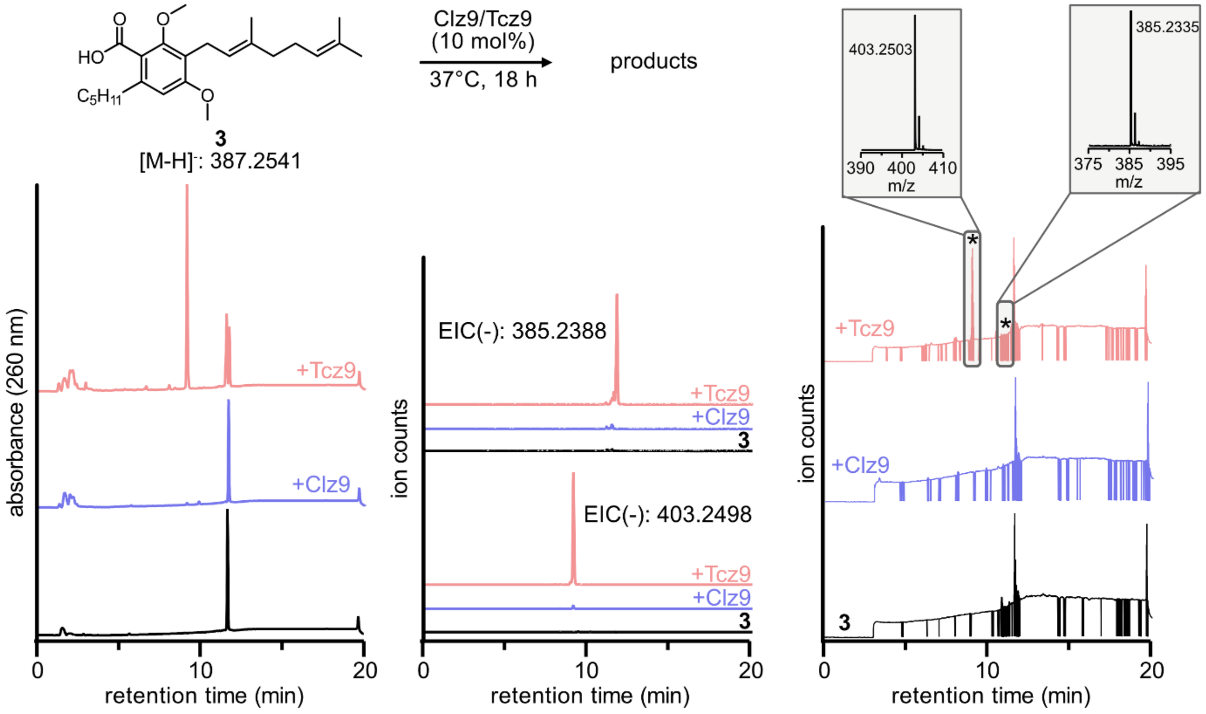


**Figure S10.** Clz9 and Tcz9 reaction with **3** to produce some products. Absorbance at 260 nm and extracted ion chromatographs of reactions with and without enzyme added are shown. TICs are also shown on the right with the inset showing exact mass for products. LC-MS was performed in negative mode.

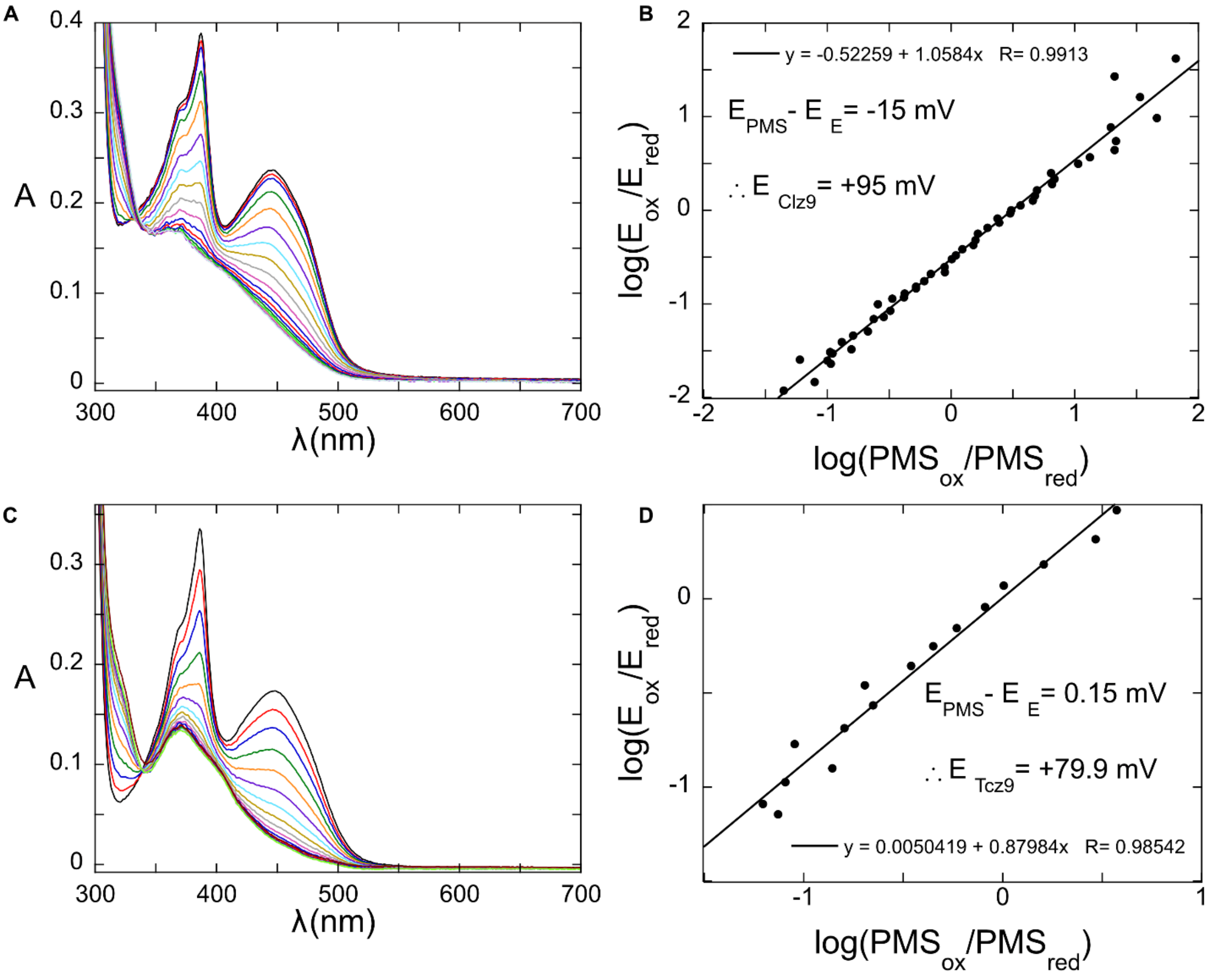


**Figure S11.** Spectroscopic determination of redox potential. Reductive titration of the flavin and corresponding Nerst equation graphs for (A-B) Clz9 and (C-D) Tcz9.

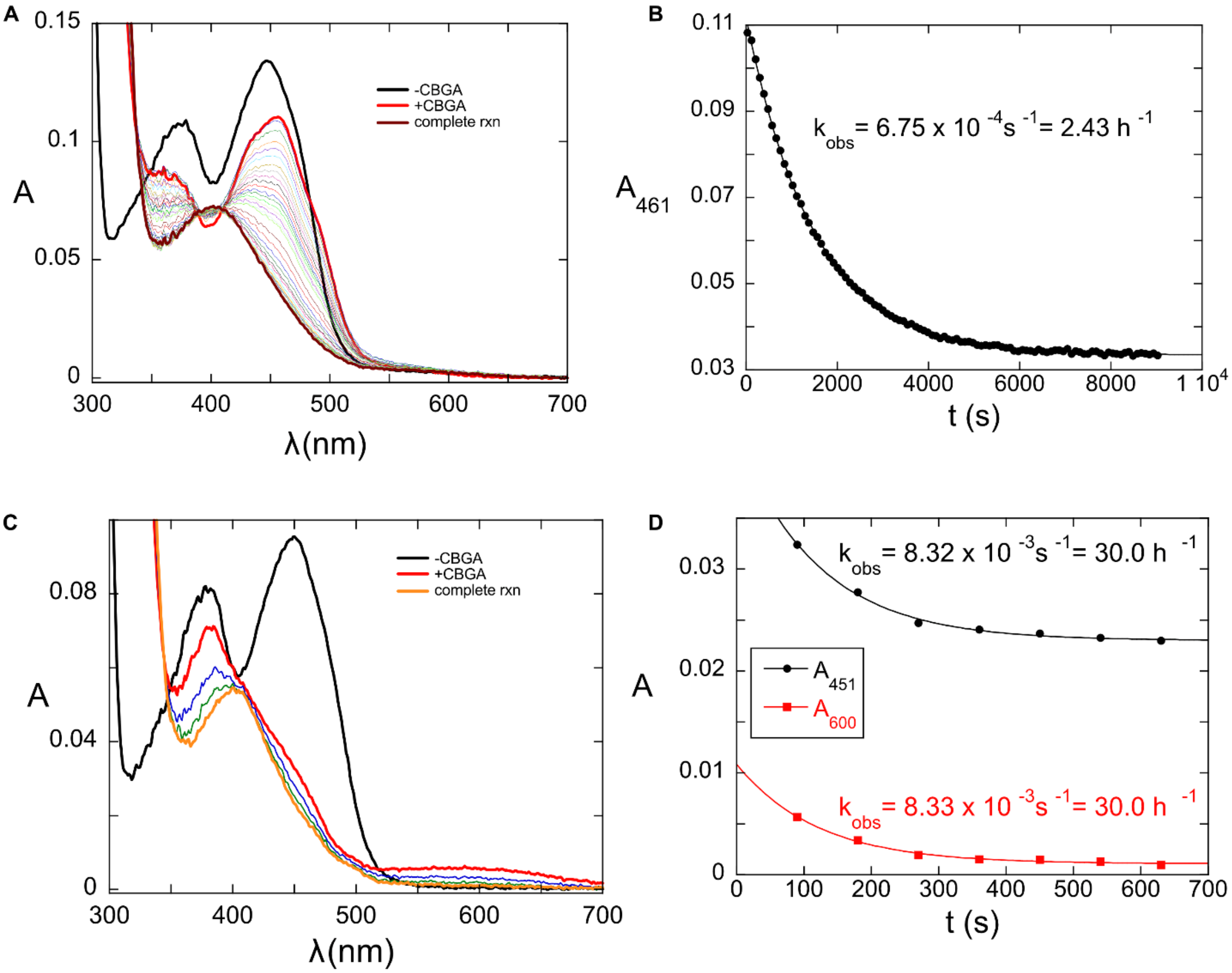


**Figure S12.** Anaerobic oxidation of **1** by Clz9 (A-B) and Tcz9 (C-D), illustrating how rapidly a single turnover takes place for Tcz9 vs. Clz9.

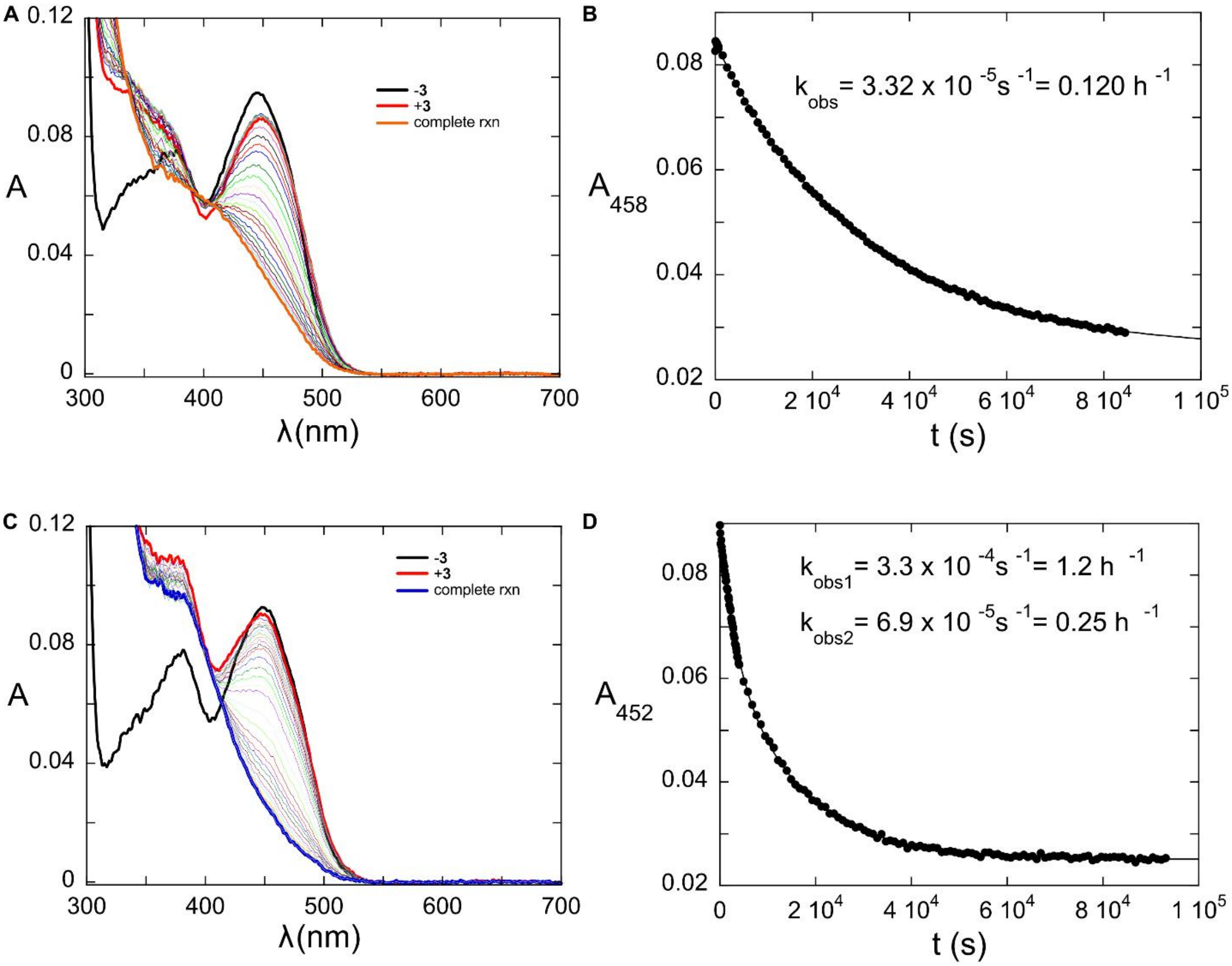


**Figure S13.** Anaerobic oxidation of **3** by Clz9 (A-B) and Tcz9 (C-D) observed spectroscopically.

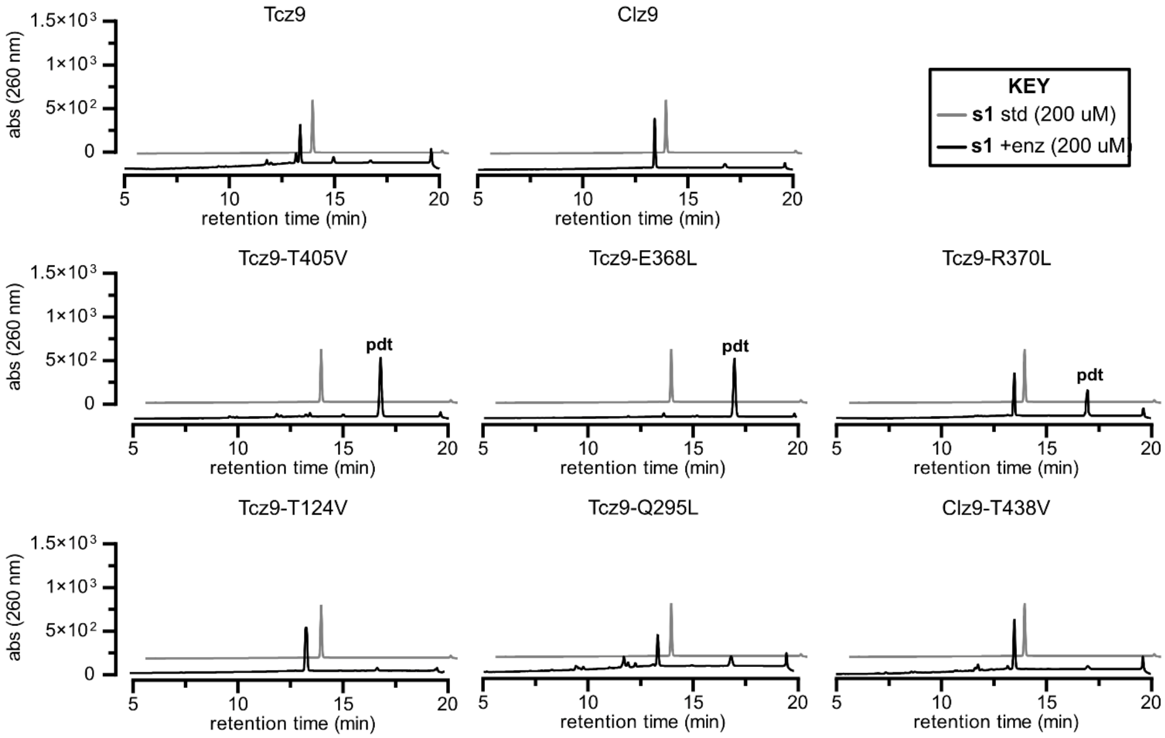


**Figure S14.** Enzyme display stark differences in reactivity with **s1**.

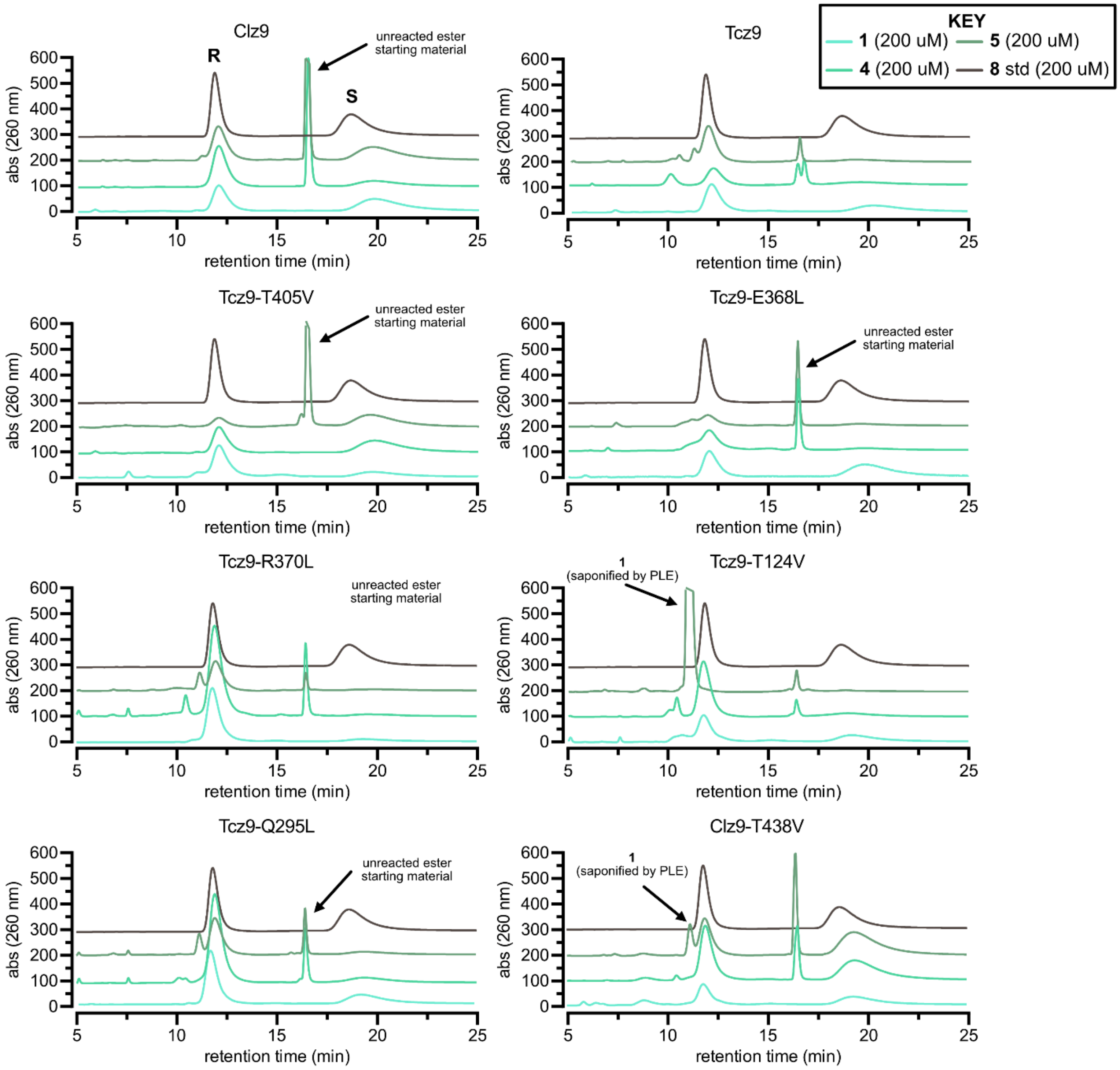


**Figure S15.** Chiral LC traces for all mutants and substrates listed in **Figure 5**.

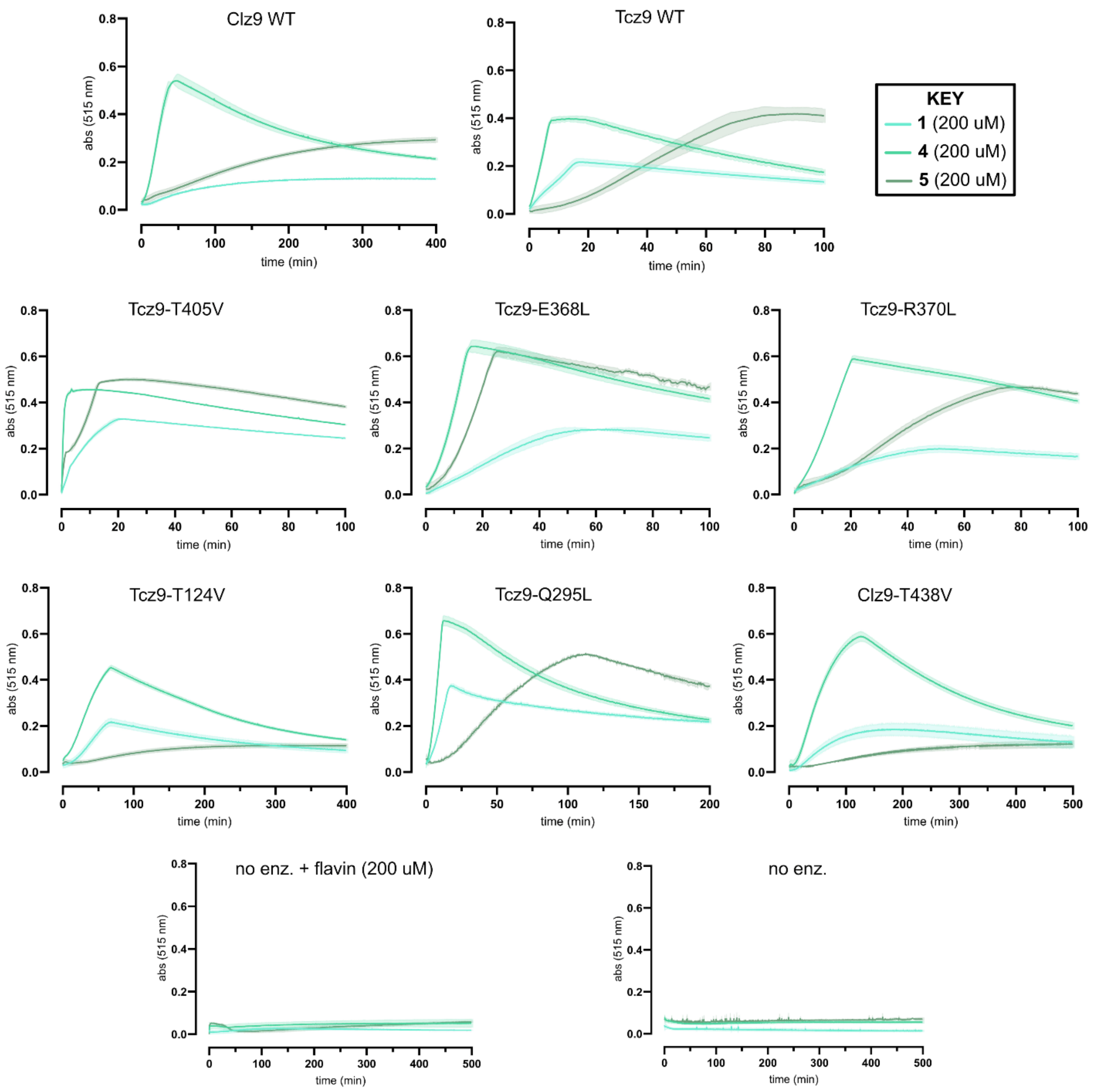


**Figure S16.** Horseradish peroxidase assay was used to calculate relative rates of turnover with **1**, **4**, and **5**.

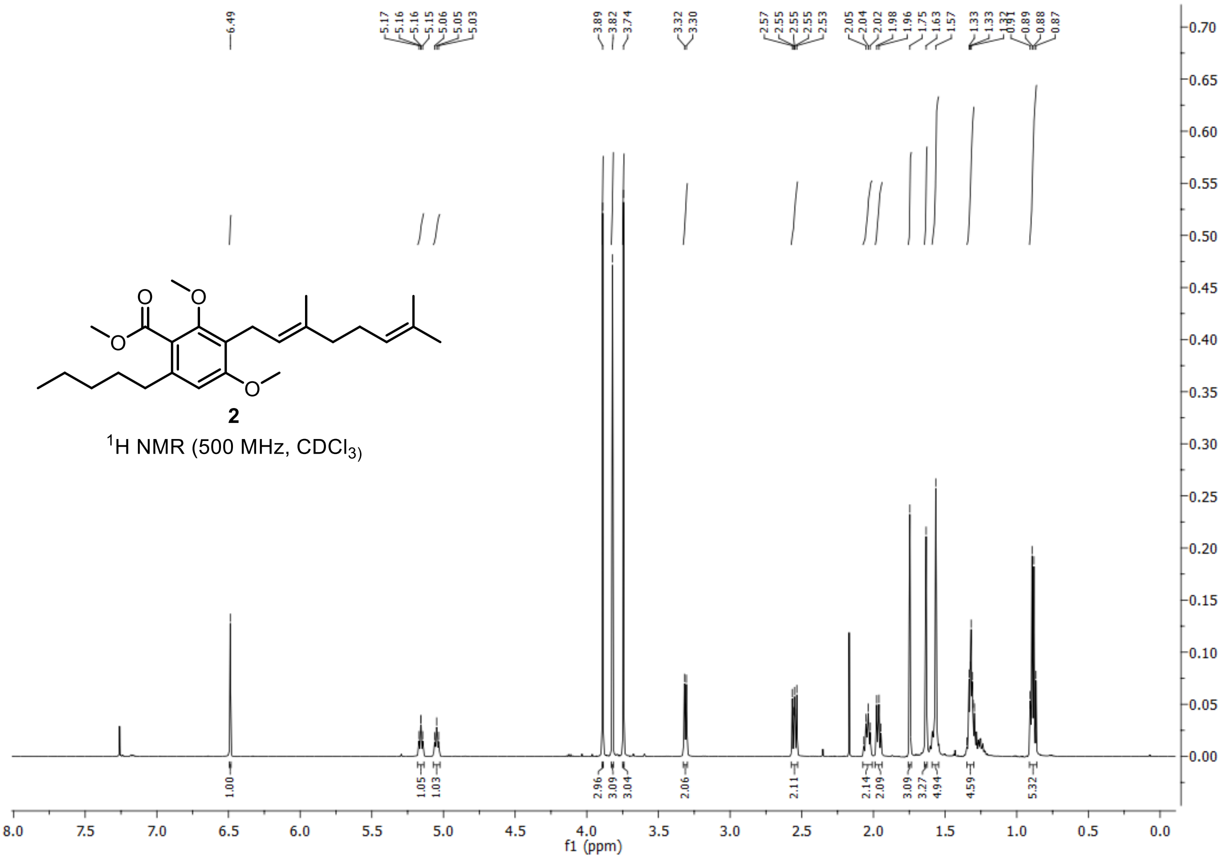
2
$^1$H NMR (500 MHz, $CDCl_3$)
f1 (ppm)

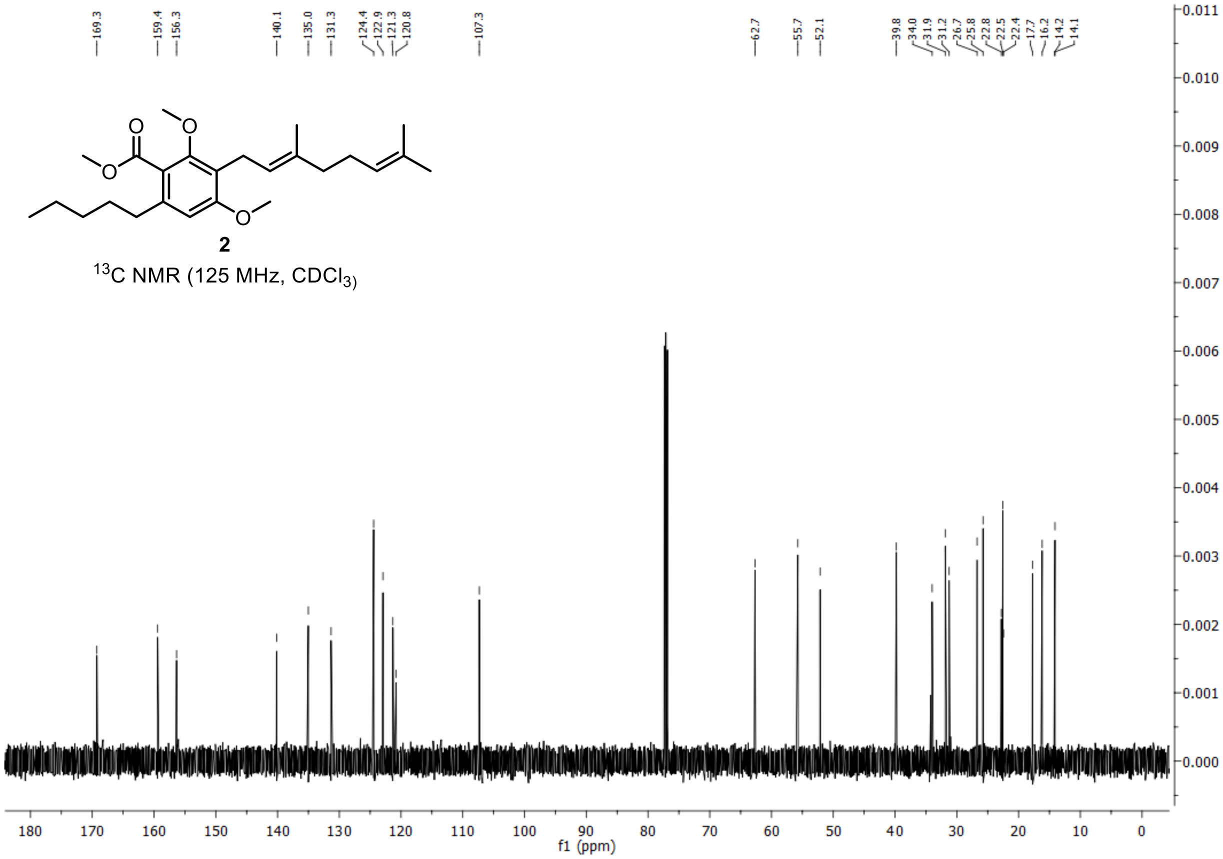
169.3
159.4
156.3
140.1
135.0
131.3
124.4
122.9
121.3
120.8
107.3
62.7
55.7
52.1
39.8
34.0
31.9
31.2
26.7
25.8
22.8
22.5
22.4
17.7
16.2
14.2
14.1
2
$^{13}$C NMR (125 MHz, $CDCl_3$)
f1 (ppm)

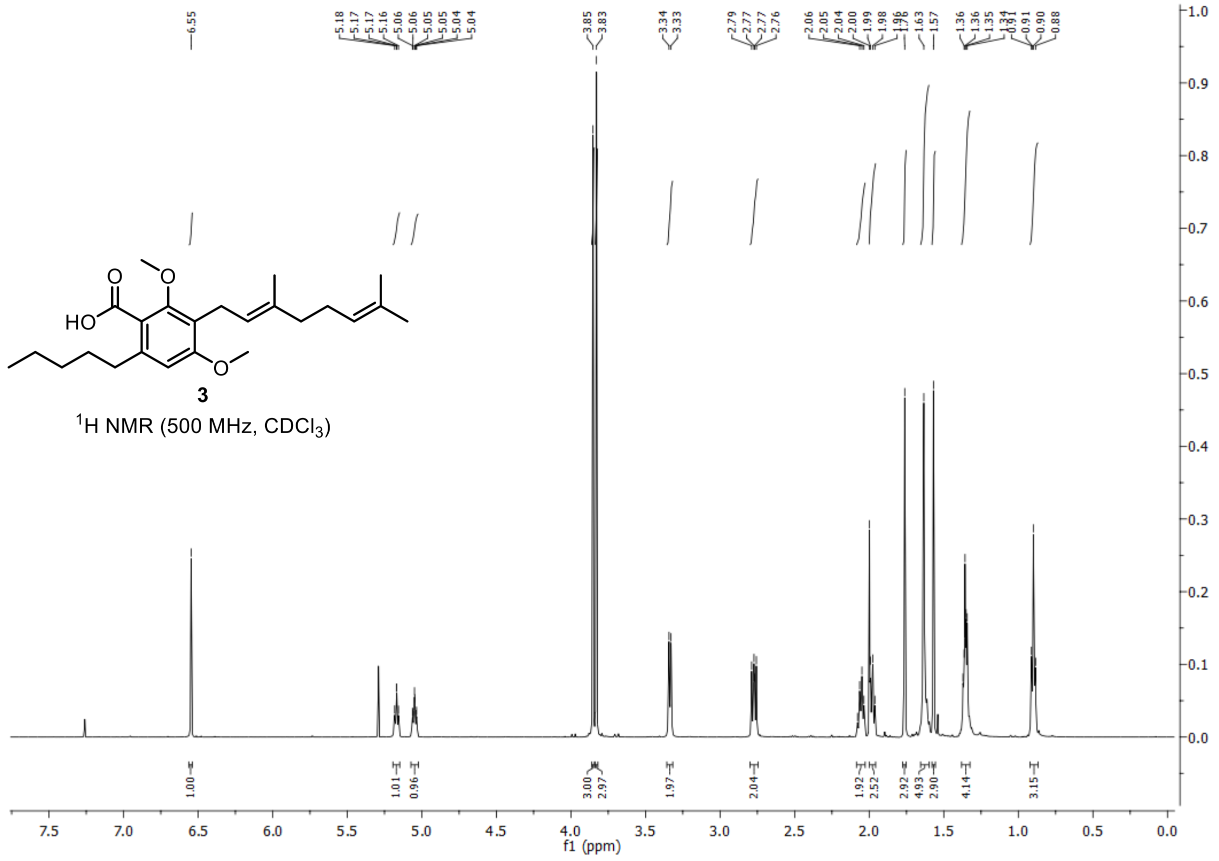
HO
3
1H NMR (500 MHz, CDCl3)
f1 (ppm)

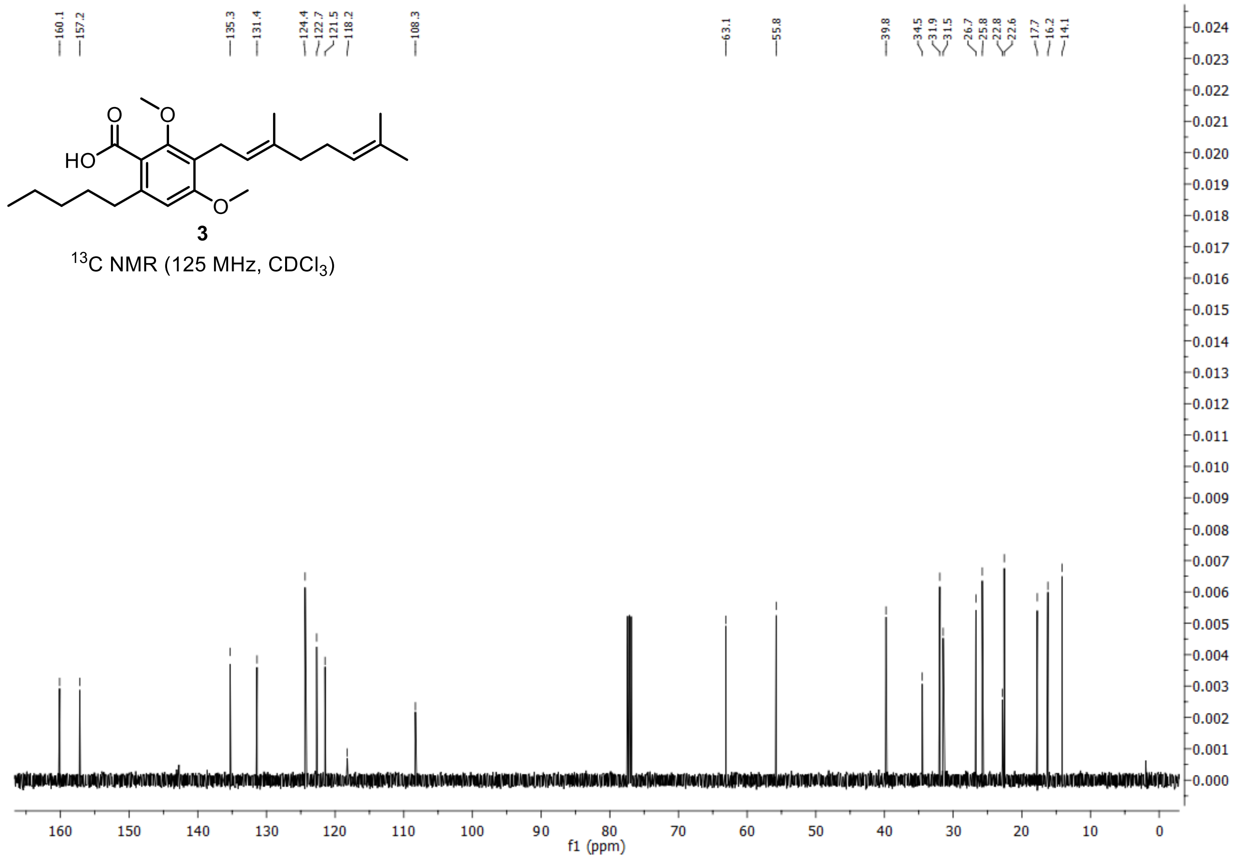


**3**

$^{13}$C NMR (125 MHz, $CDCl_3$)

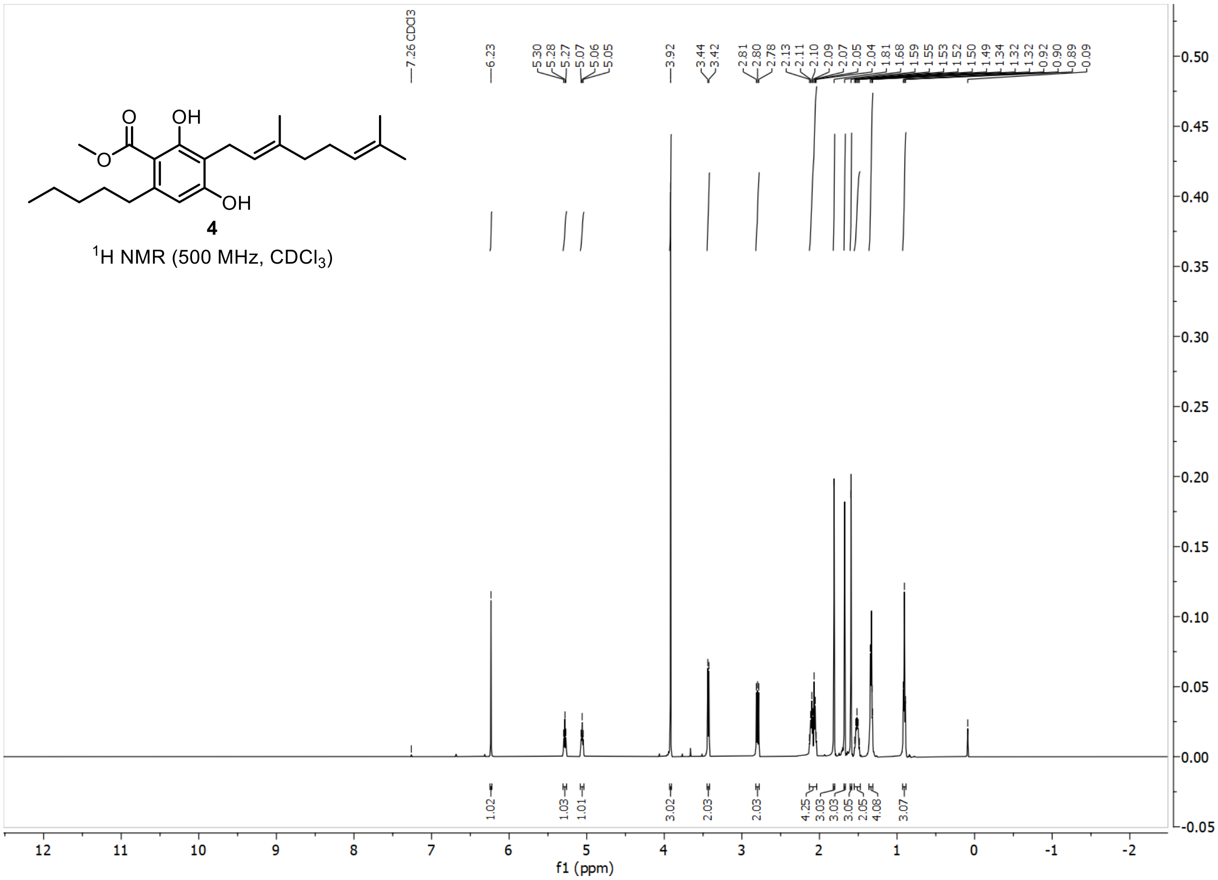

**4**

$^1$H NMR (500 MHz, $CDCl_3$)

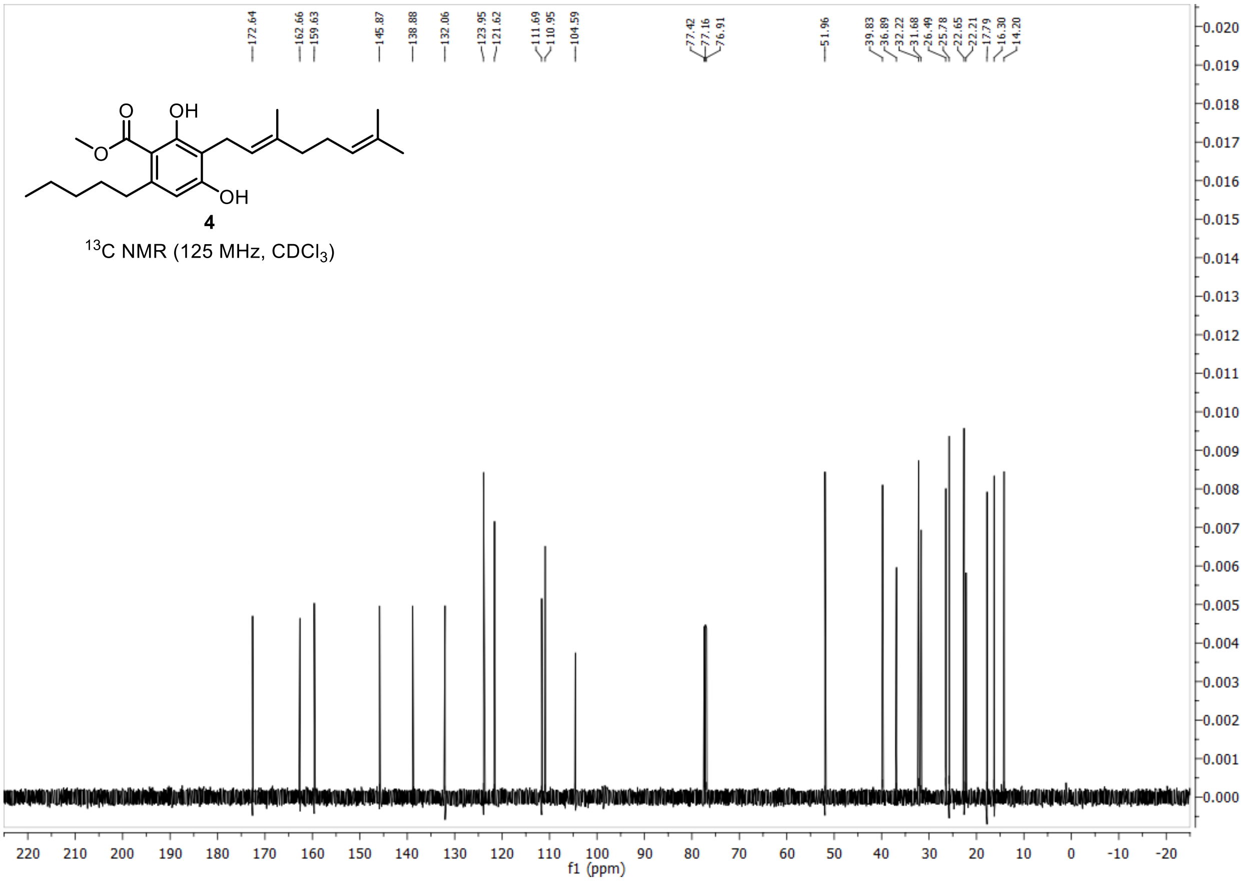


$^{13}C$ NMR (125 MHz, $CDCl_3$)

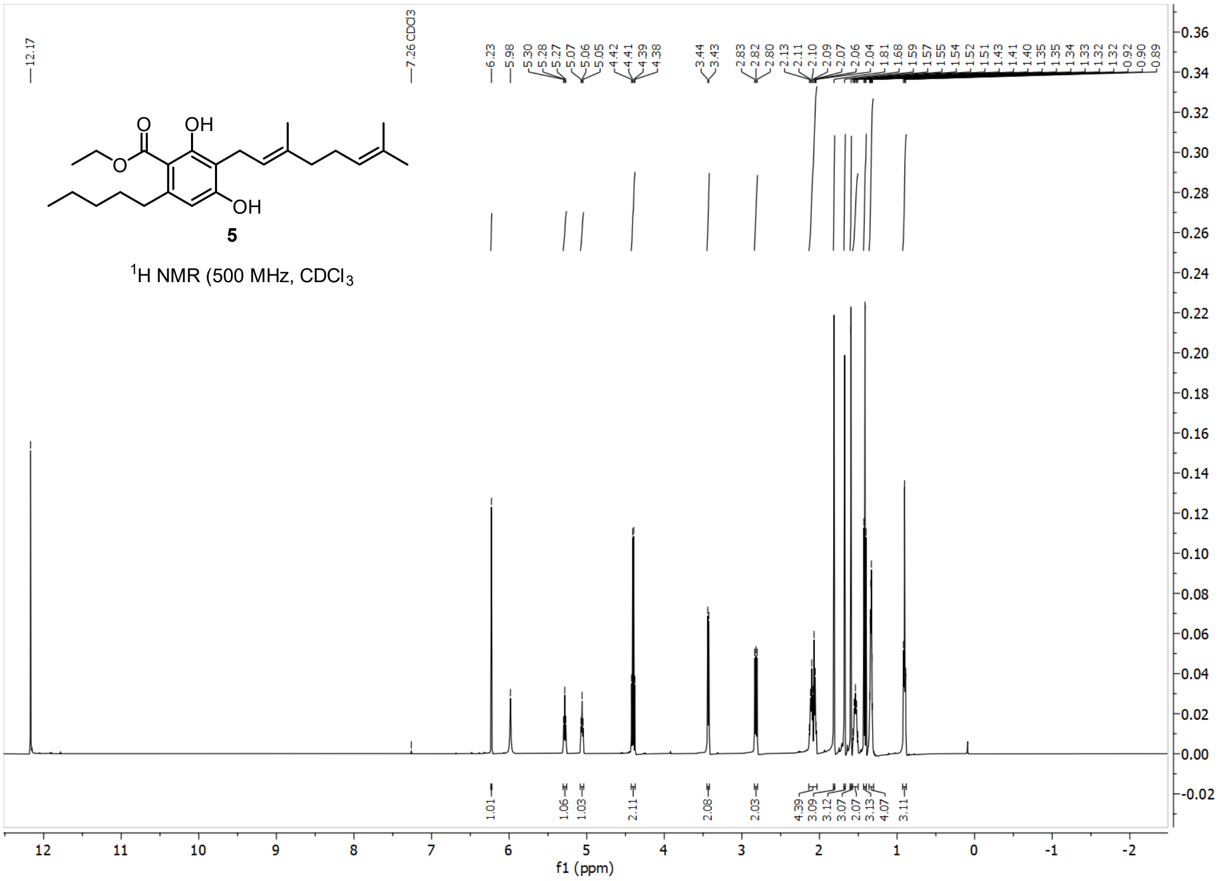

OH
O
O
OH
5
1H NMR (500 MHz, CDCl3
12.17
7.26 CDCl3
f1 (ppm)

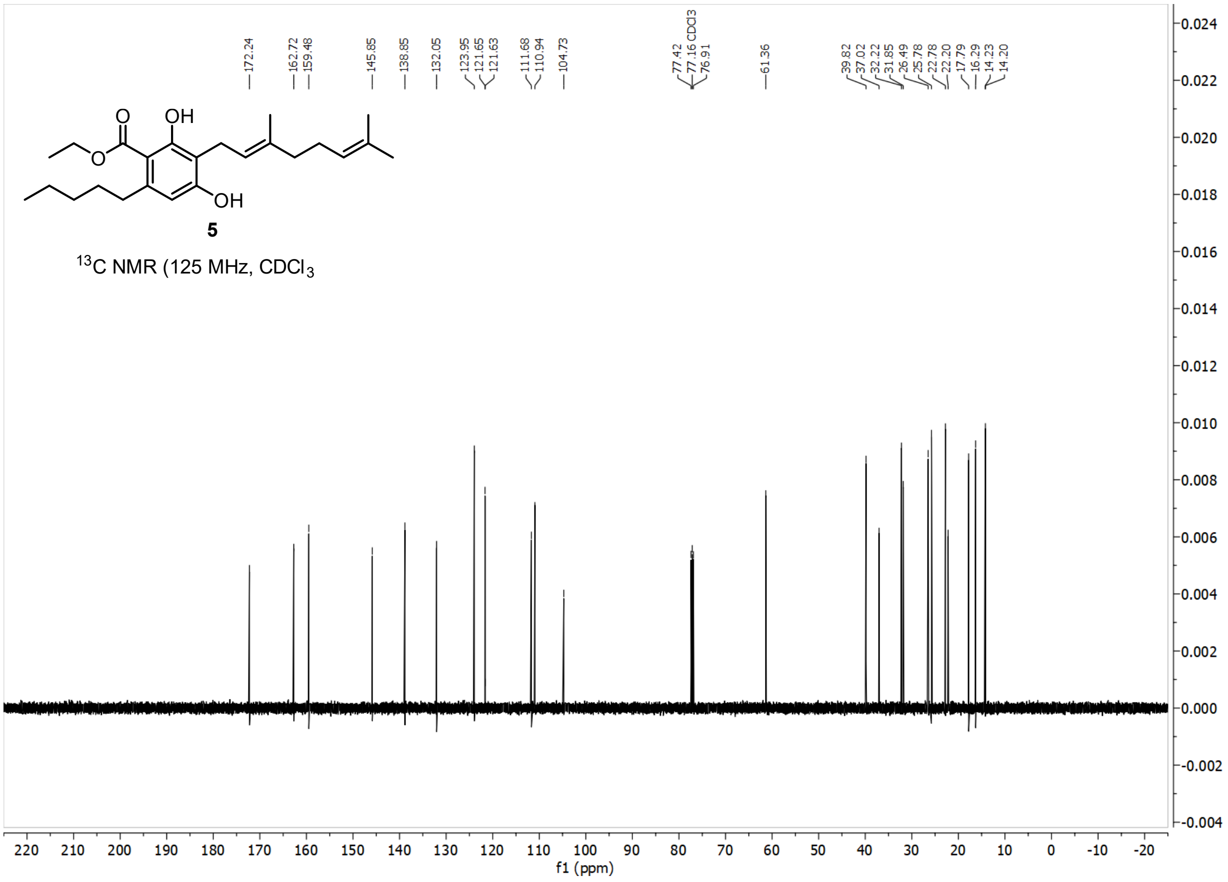

172.24
162.72
159.48
145.85
138.85
132.05
123.95
121.65
121.63
111.68
110.94
104.73
77.42
77.16 CDCl3
76.91
61.36
39.82
37.02
32.22
31.85
26.49
25.78
22.78
22.20
17.79
16.29
14.23
14.20
O
OH
O
OH
5
$^{13}$C NMR (125 MHz, $CDCl_3$
f1 (ppm)

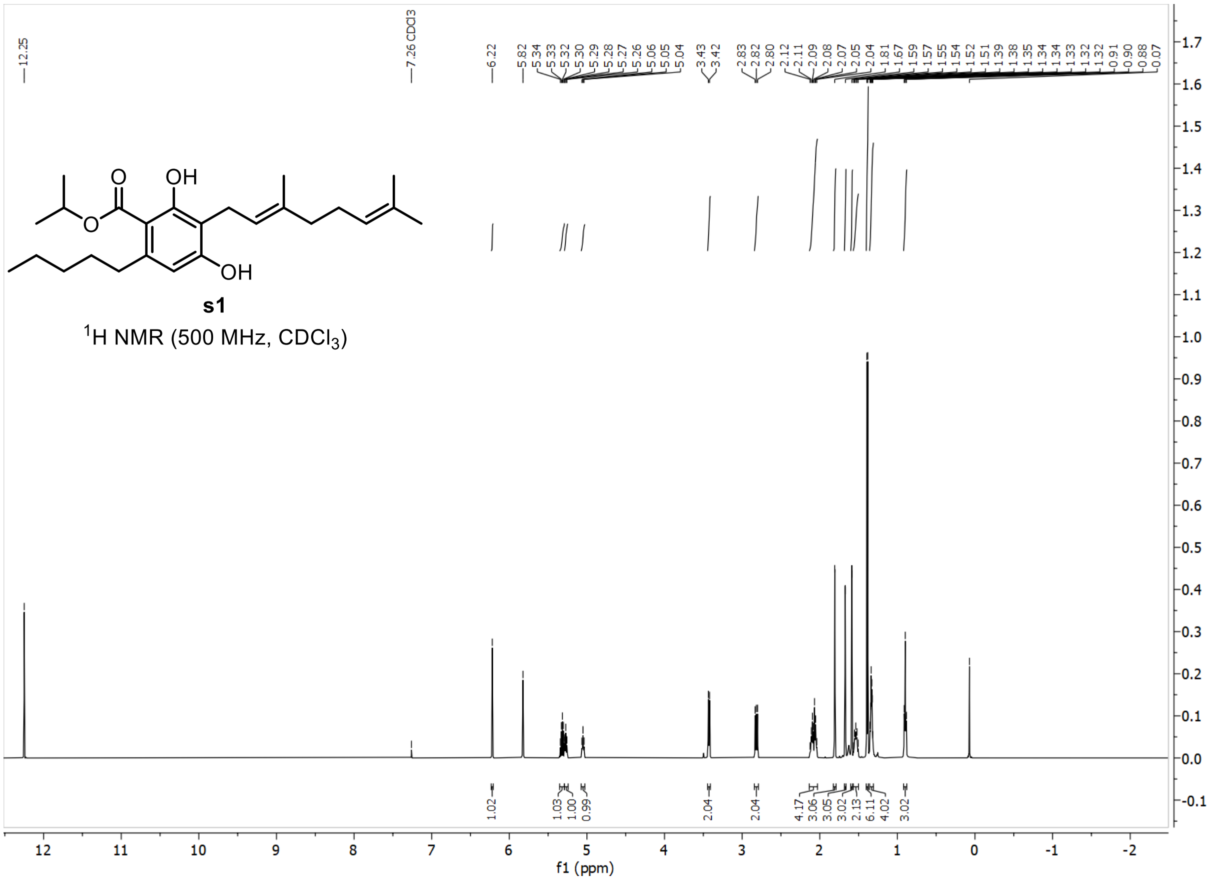

12.25
7.26 CDCl3
6.22
5.82
5.34
5.33
5.32
5.30
5.29
5.28
5.27
5.26
5.06
5.05
5.04
3.43
3.42
2.83
2.82
2.80
2.12
2.11
2.09
2.08
2.07
2.05
2.04
1.81
1.67
1.59
1.57
1.55
1.54
1.52
1.51
1.39
1.38
1.35
1.34
1.34
1.33
1.32
1.32
0.91
0.90
0.88
0.07
OH
O
O
OH
s1
1H NMR (500 MHz, CDCl3)
1.02
1.03
1.00
0.99
2.04
2.04
4.17
3.06
3.05
3.02
2.13
6.11
4.02
3.02
f1 (ppm)

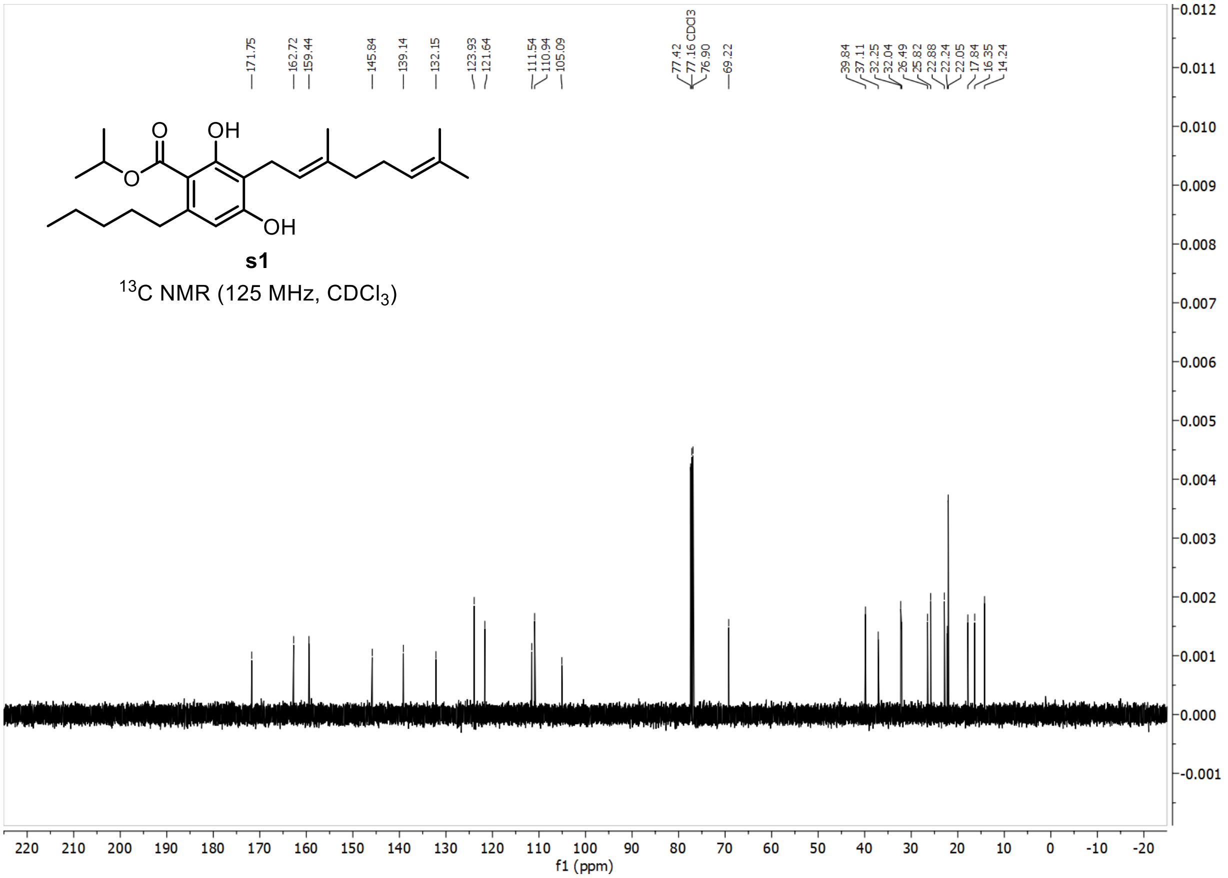
171.75
162.72
159.44
145.84
139.14
132.15
123.93
121.64
111.54
110.94
105.09
77.42
77.16 CDCl3
76.90
69.22
39.84
37.11
32.25
32.04
26.49
25.82
22.88
22.24
22.05
17.84
16.35
14.24
OH
O
O
OH
s1
13C NMR (125 MHz, CDCl3)
f1 (ppm)
